# Dietary palmitic acid induces innate immune memory via ceramide production that enhances severity of acute septic shock and clearance of infection

**DOI:** 10.1101/2021.06.15.448579

**Authors:** AL Seufert, JW Hickman, SK Traxler, RM Peterson, SJ Lashley, N Shulzhenko, RJ Napier, BA Napier

## Abstract

Trained immunity is an innate immune memory response that is induced by primary microbial or sterile stimuli that sensitizes monocytes and macrophages to a secondary pathogenic challenge, reprogramming the host response to infection and inflammatory disease. Nutritional components, such as dietary fatty acids, can act as inflammatory stimuli, but it is unknown if they can act as the primary stimuli in the context of innate immune memory. Here we find mice fed diets enriched in saturated fatty acids (SFAs) confer a hyper-inflammatory response to systemic lipopolysaccharide (LPS) and increased mortality, independent of diet-induced microbiome and glycemic modulation. *Ex vivo*, we show monocytes and splenocytes from mice fed enriched SFAs do not have altered baseline inflammation, but enhanced responses to a secondary inflammatory challenge. Lipidomics identified enhanced free palmitic acid (PA) and PA-associated lipids in SFA-fed mice serum. We found pre-treatment with physiologically relevant concentrations of PA alone reprograms macrophages to induce a hyper-inflammatory response to secondary challenge with LPS. This response was found to be dependent on the synthesis of ceramide, and reversible when treated with oleic acid, a mono-unsaturated FA that depletes intracellular ceramide. *In vivo*, we found systemic PA confers enhanced inflammation and mortality during an acute septic response to systemic LPS, which was not reversible for up to 7 days post-PA-exposure. While PA-treatment is harmful for acute septic shock outcome, we find PA exposure enhanced clearance of *Candida albicans* in RAG^-/-^ mice. These are the first data to implicate enriched dietary SFAs, and specifically PA, in the induction of long-lived innate immune memory that is detrimental during an acute septic response, but beneficial for clearance of pathogens.

## Introduction

Historically, immune memory has been defined as a trait limited to the adaptive immune system, however it is now well established that innate immune cells have the capacity for metabolic, epigenetic, and functional reprogramming that leads to long-lasting increases in host resistance to infection (*1-4*). Specifically, trained immunity is an adaptation of innate host defense in vertebrates and invertebrates that results from exposure to a primary inflammatory stimulus and leads to a faster and greater response to a secondary challenge. Unlike adaptive memory responses, trained immunity does not require genome rearrangements, B and T lymphocytes, and receptors that recognize antigens (*1-4*). Further, trained immunity has been documented in organisms that lack canonical adaptive immune responses, such as plants and invertebrates, suggesting this is a primitive immune memory system that is conserved throughout vertebrates and invertebrates (*5*).

The Bacillus Calmette-Guérin (BCG) vaccine and yeast β-glucans are canonical inducers of trained immunity in humans, and stimulate long-lasting metabolic and epigenetic reprogramming of myeloid-lineage cells resulting in hyperresponsiveness upon restimulation with heterologous or homologous inflammatory stimuli. This innate immune memory is heritable (*6*) and can last up to months in humans and mice (*7*); thus, likely evolved to provide non-specific protection from secondary infections. Most recently, it was described that countries with higher rates of BCG vaccine at birth had fewer coronavirus disease 2019 (COVID-19) cases (*8*) making this immunological phenomenon extremely relevant. Importantly, it is easily ascertained that inflammatory hyperresponsiveness could be deleterious in the context of diseases where more inflammation can lead to greater pathology (ex: acute septic shock, autoimmune disorders, and allergies). Thus, trained immunity can be regarded as a double-edged sword – providing increased resistance to infection but exacerbating inflammatory diseases. Consequently, identifying novel inducers of trained immunity will provide clinically relevant insight into harnessing innate immune cells to attain long-term therapeutic benefits in a range of infections and inflammatory diseases.

Typically, the primary inflammatory stimulus that initiates trained immunity are danger- or pathogen-associated molecular patterns (DAMPs; PAMPs); however, recent publications have shown dietary components (*e*.*g*., β-glucan found in mushrooms, baker’s and brewer’s yeast, wheat and oats, and unknown components of bovine milk) can induce trained innate immune memory and alter inflammatory disease outcome (*9, 10*). These data contribute to the growing evidence supporting the multifaceted immunoregulatory role of diet.

Currently, Westernized nations are increasingly dependent on diets enriched in saturated fatty acids (SFAs) (*11-13*), which have been shown to mimic PAMP effects on inflammatory cells, regulate innate immune cell function, and alter outcomes of inflammatory disease and infection (*14-17*). Specifically, we have shown the Western diet (WD), a diet enriched in sucrose and SFAs, correlates with increased disease severity and mortality in response to systemic LPS, independent of the diet-dependent microbiota, demonstrating the possibility that the dietary components of this diet may be driving the hyperresponsiveness to LPS (*18*). Currently, it is unknown if enriched dietary SFAs mediate trained immunity.

Our work presented herein identifies a diet enriched in SFAs, and not sucrose, confers an increased systemic response to LPS independent of microbiome or glycolytic regulation during disease. A lipidomic analysis of blood fat composition after high SFA-feeding revealed a significant increase of free palmitic acid (PA: C16:0) and fatty acid complexes containing PA. We find exposure to palmitic acid (PA; C16:0), the predominant SFA found in Westernized diets, does not affect baseline inflammation but enhances systemic response to microbial ligands and resistance to infection. We identify a novel role of PA-dependent intracellular ceramide required for this enhanced response, and show intervention with oleic acid, a mono-unsaturated fatty acid that depletes PA-dependent ceramide, can reverse these phenotypes. These data are consistent with the current knowledge that PA and ceramide are immunomodulatory molecules, and build on these by highlighting a previously unidentified role of PA in driving long-lived trained immunity.

## Results

### Diets enriched in saturated fatty acids increase endotoxemia severity and mortality

To examine the immune effects of chronic exposure to diets enriched in SFAs on lipopolysaccharide (LPS)-induced acute septic shock, we fed mice either a WD (enriched in SFAs and sucrose), a ketogenic diet (KD; enriched in SFAs and low-carbohydrate), or standard chow (SC; low in SFAs and sucrose), for 2 weeks (wk) (Table S1). We defined 2 wk of feeding as chronic exposure, because this is correlated with WD- or KD-dependent microbiome changes, and confers metaflammation in WD mice (*18*), sustained altered blood glucose levels (Fig S1A), and elevated levels of ketones in the urine and blood (Fig S1B, C) (*19*). We then induced acute septic shock by a single intraperitoneal (i.p.) injection of LPS. We measured hypothermia as a measure of disease severity and survival to determine outcome (*18, 20, 21*). WD- and KD-fed mice showed significant and prolonged hypothermia, starting at 10 hours (h) post-injection (p.i.), compared to the SC-fed mice (Fig 1A). In accordance with these findings, WD- and KD-fed mice displayed 100% mortality by 26 h p.i. compared to 100% survival of SC-fed mice (Fig 1B). Hypoglycemia is a known driver of acute septic mortality, and each of these diets has varying levels of sugars and carbohydrates (Table S1) (*22, 23*). However, mice in all diet groups displayed similar levels of LPS-induced hypoglycemia during disease (Fig S1D), indicating that potential effects of diet on blood glucose were not a driver of enhanced disease severity.

**Fig. 1:**
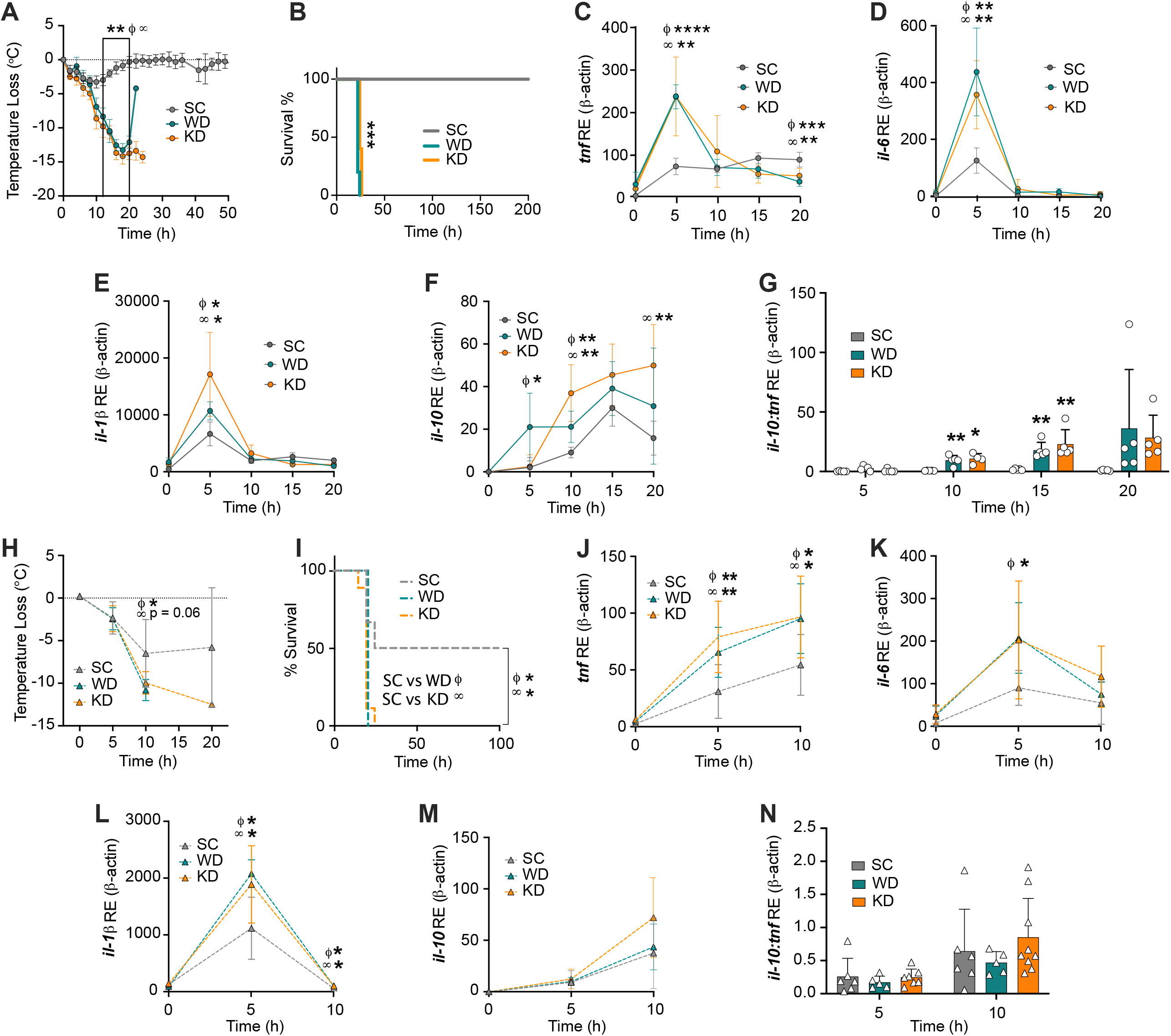
Diets enriched in SFAs lead to enhanced acute septic shock severity and altered systemic inflammatory profiles, independent of microbiome. Age-matched (4-6 wk) conventional mice were fed SC, WD, or KD for 2 wk and injected i.p. with 6 mg/kg of LPS. **(A)** Temperature loss and **(B)** survival were monitored every 2 h. At indicated times 10-20 μL of blood was drawn via the tail vein, RNA was collected, and samples were assessed for expression of **(C)** *tnf*, **(D)** *il-6*, **(E)** *il-1β*, and **(F)** *il-10* via qRT-PCR. **(G)** *il-10:tnf* ratio was calculated for 5, 10, 15, and 20 hours p.i. with LPS. Next, 19-23 wk old female and 14-23 wk old male germ-free C57BL/6 mice were fed SC, WD, or KD for 2 wk and injected i.p. with 50 mg/kg of LPS. **(H)** Temperature loss and **I** survival were monitored every 5 h p.i. **(J-N)** At indicated times, 10-20 μL of blood was drawn via the tail vein, RNA was collected, and samples were assessed for expression of **(J)** *tnf*, **(K)** *il-6*, **(L)** *il-1β*, and **(M)** *il-10* via qRT-PCR. **(N)** *il-10:tnf* ratio was calculated for 5 and 10 h p.i. with LPS. SC, n=6; WD, n= 5, and KD, n= 9. For **A, C-G, H**, and **J-N** a Mann Whitney test was used for pairwise comparisons. For **B** and **I** a log-rank Mantel-Cox test was used for survival curve comparison. For **A-N**, data are representative of 1 experiment, n=5-9 mice per diet group. For **A, C-G, H, and J-N** a Mann Whitney test was used for pairwise comparisons. For all panels, \**p*< 0.05; \*\**p* < 0.01; \*\*\**p*< 0.001. For **c-e**, Φ symbols indicate WD significance and ∞ symbols indicate KD significance. Error bars shown mean ± SD.

Considering mice fed KD experience a shift towards nutritional ketosis, we wanted to understand if our phenotype was dependent on nutritional ketosis. 1,3-butanediol is a compound that induces ketosis by enhancing levels of the ketone β-hydroxybutyrate in the blood (*19*). Mice were fed for 2 wk with KD, SC supplemented with saccharine and 1,3-butanediol (SC + BD) or SC-fed with the saccharine vehicle solution (SC + Veh). We next injected LPS i.p. and found KD-fed mice showed significantly greater hypothermia, and increased mortality, compared to SC + BD and SC + Veh (Fig S1E, F). Though short-lived, when compared to SC + Veh, the SC + BD mice did confer an increase in hypothermia, suggesting that nutritional ketosis may play a minor role in KD-dependent susceptibility to acute septic shock (Fig S1E, F). Together these data suggest that diets enriched in SFAs promote enhanced acute septic shock severity and this is independent of diet-dependent hypoglycemic shock or nutritional ketosis.

### Diets enriched in SFAs induce a hyper-inflammatory response to LPS and increased immunoparalysis

LPS-induced acute septic shock mortality results exclusively from a systemic inflammatory response, characterized by an acute increase in circulating inflammatory cytokine levels (ex: TNF, IL-6, and IL-1β) from splenocytes and myeloid derived innate immune cells (*24-27*). Additionally, pre-treatment of myeloid-derived cells with dietary SFAs has been shown to enhance inflammatory pathways in response to microbial ligands (*28, 29*). Considering this, we hypothesized that exposure to enriched systemic dietary SFAs in WD- and KD-fed mice would enhance the inflammatory response to systemic LPS during the acute septic response. Five-hours p.i., mice fed all diets showed induction of *tnf, il-6*, and *il-1β* expression in the blood (Fig 1C-E). However, WD- and KD-fed mice experienced significantly higher expression of *tnf, il-6*, and *il-1β* in the blood at 5 h p.i., compared with SC-fed mice (Fig 1C-E), indicating that diets enriched in SFAs are associated with a hyper-inflammatory response to LPS.

Importantly, septic patients often present with two immune phases: an initial amplification of inflammation, followed-by or concurrent-with an induction of immune suppression (immunoparalysis), that can be measured by a systemic increase in the anti-inflammatory cytokine IL-10 (*30, 31*). Further, in septic patients, a high IL-10:TNF ratio equates with the clinical immunoparalytic phase and correlates with poorer sepsis outcomes (*32, 33*). Interestingly, we found there was significantly increased *il-10* expression in WD-fed mice at 5 and 10 h and KD-fed mice at 10 and 20 h p.i., compared to SC-fed mice (Fig 1F), and WD- and KD-fed mice had significantly higher *il-10:tnf* ratios at 10 and 15 h compared to SC-fed mice (Fig 1G). These data conclude that mice exposed to diets enriched in SFAs show an initial hyper-inflammatory response to LPS, followed by an increased immunoparalytic phenotype, which correlates with enhanced disease severity, similar to what is seen in the clinic.

### Diets enriched in SFAs drive enhanced responses to systemic LPS independent of the microbiome

We have previously shown that WD-fed mice experience increased acute septic shock severity and mortality, independent of the microbiome (*18*). In order to confirm the increases in disease severity that correlated with KD were also independent of KD-associated microbiome changes, we used a germ free (GF) mouse model. GF mice were fed SC, WD, and KD for 2 wk followed by injection with 50 mg/kg of LPS, our previously established LD_50_ (*18*). As we saw in the conventional mice, at 10 h p.i. WD- and KD-fed GF mice showed enhanced hypothermia and mortality, compared to SC-fed GF mice (Fig 1H, I). These data show, similar to WD-fed mice, KD-associated increase in acute septic shock severity and mortality is independent of the microbiome.

Additionally, to confirm that the hyper-inflammatory response to systemic LPS was independent of the WD- and KD-dependent microbiome, we measured systemic inflammation during endotoxemia via the expression of *tnf, il-6*, and *il-1β* in the blood at 0-10 h p.i. We found, WD- and KD-fed GF mice displayed significantly enhanced expression of *tnf* and *il-1β* at 5-10 h, and WD-fed mice showed enhanced expression of *il-6* at 5 h, compared to SC-fed GF mice (Fig 1J-L). Interestingly, *il-10* expression and the *il-10:tnf* ratio were not significantly different throughout all diets, suggesting the SFA-dependent enhanced immunoparalytic phenotype is dependent on the diet-associated microbiomes in WD- and KD-fed mice (Fig 1M, N). These data demonstrate that the early hyper-inflammatory response, but not the late immunoparalytic response, to LPS associated with enriched dietary SFAs is independent of the diet-dependent microbiota.

### Monocytes and splenocytes from KD-fed mice show a hyper-inflammatory response to LPS

As mentioned previously, monocytes and splenocytes are necessary for induction of systemic inflammatory cytokines during LPS-initiated acute septic shock (*26, 27*). Here we see feeding a diet enriched only in SFAs (KD) leads to enhanced expression of inflammatory cytokines in the blood during acute septic shock; however, it remains unclear if the KD induces *in vivo* reprogramming of monocytes and splenocytes leading to an enhanced response to LPS. Thus, we sought out to determine if the chronic exposure to KD alters the response of monocytes and splenocytes to LPS *ex vivo*. First, we fed mice SC or KD for 2 wk (chronic exposure), isolated bone marrow monocytes (BMMs) and determined baseline inflammation of monocytes via expression of inflammatory cytokines. We found that prior to *ex vivo* LPS stimulation (0 h), BMMs isolated from mice chronically exposed to SC- or KD showed no significant difference in *tnf* expression, and, interestingly, *il-6* expression was significantly decreased in BMMs from KD-fed mice (Fig 2A). However, when BMMs were stimulated with LPS for 2 h *ex vivo*, those from KD-fed mice showed significantly higher expression of *tnf* and *il-6* (Fig 2A). Similarly, we isolated splenocytes from SC- and KD-fed mice and found no difference between homeostatic inflammation of splenocytes between diets (0 h), but an enhanced production of *tnf* in the splenocytes of KD-fed mice challenged with LPS (2 h), compared to splenocytes from SC-fed mice (Fig 2B). Together, these data suggest that BMMs and splenocytes from KD-fed mice are not more inflammatory in a healthy mouse, but confer a hyper-inflammatory response to a secondary inflammatory stimulus, suggesting diets enriched in SFAs are inducing functional reprogramming of immune cells *in vivo*.

**Fig. 2:**
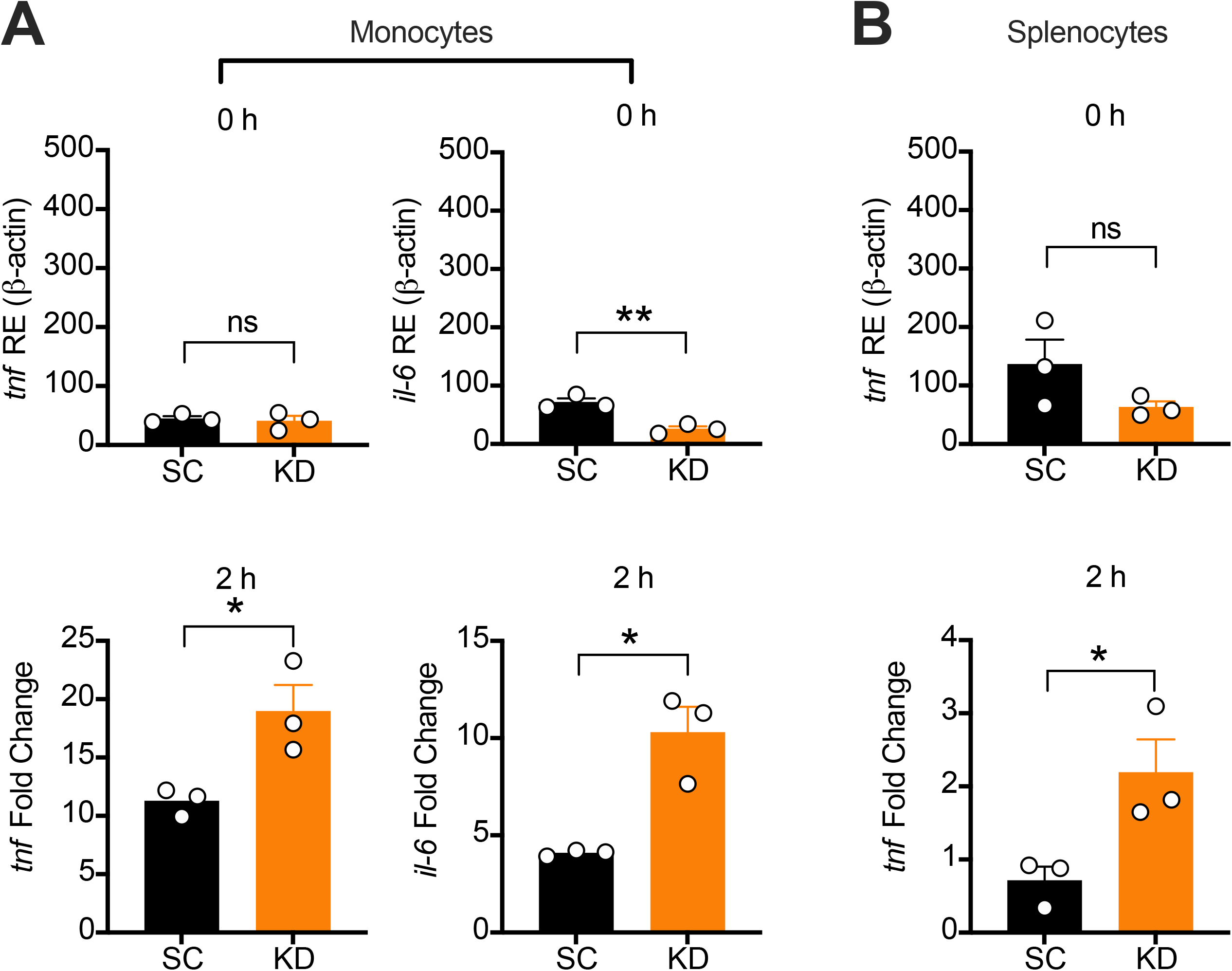
Monocytes and splenocytes from KD-fed mice show a hyper-inflammatory response to LPS *ex vivo*. Age-matched (4-6 wk) mice fed SC or KD for 2 weeks. Monocytes were isolated from the femurs and tibias of mice and plated at 2×10^^6^ cells/mL. RNA was extracted from (**A)** untreated monocytes (0 h) or monocytes treated with or without LPS (10 ng/mL) for 2 h. Expression of *tnf* and *il-6* was analyzed via qRT-PCR. Splenocytes were isolated and plated at 1×10^^6^ cells/mL. RNA was isolated from **(B)** untreated splenocytes (0 h) or splenocytes treated with or without LPS (10 ng/mL) for 2 h. Expression of *tnf* and *il-6* was analyzed via qRT-PCR. *n* = 5 mice/group in each representative experiment. A student’s t-test was used for statistical significance. For all panels, * *p* < 0.05; ** *p* < 0.01; *** *p* < 0.001; **** *p* < 0.0001.

### Palmitic acid (PA) and PA-associated fatty acids are enriched in the blood of KD-fed mice

We next wanted to identify target dietary SFAs enriched in the blood of mice that may be altering the systemic inflammatory response to LPS. It is known that the SFAs consumed in the diet determine the SFA profiles in the blood (*34-36*). Considering this, we used mass spectrometry lipidomics to create diet-dependent profiles of circulating fatty acids in SC- and KD-fed mice (*37*). Mice were fed SC or KD for 2 wk, then serum samples were collected and analyzed using qualitative tandem liquid chromatography quadrupole time of flight mass spectrometry (LC-QToF MS/MS). We used principal component analysis (PCA) to visualize how samples within each data set clustered together according to diet, and how those clusters varied relative to one another in abundance levels of free fatty acids (FFA), triacylglycerols (TAG), and phosphatidylcholines (PC). For all three groups of FAs, individual mice grouped with members of the same diet represented by a 95% confidence ellipse with no overlap between SC- and KD-fed groups (Fig 3A-C). These data indicate that 2 wk of KD feeding is sufficient to significantly alter circulating FFAs, TAGs, and PCs, and that SC- and KD-fed mice display unique lipid blood profiles. Similarly, the relative abundance of sphingolipids (SG) in SC- and KD-fed mice displayed unique diet-dependent profiles with no overlapping clusters (Fig S2A). Though the independent role of each FFA, TAG, PC, and SG species has not been clinically defined, each are classes of lipids that when accumulated is associated with metabolic diseases, which have been shown to enhance susceptibility to sepsis and exacerbate inflammatory disease (*16, 38-40*).

**Fig. 3:**
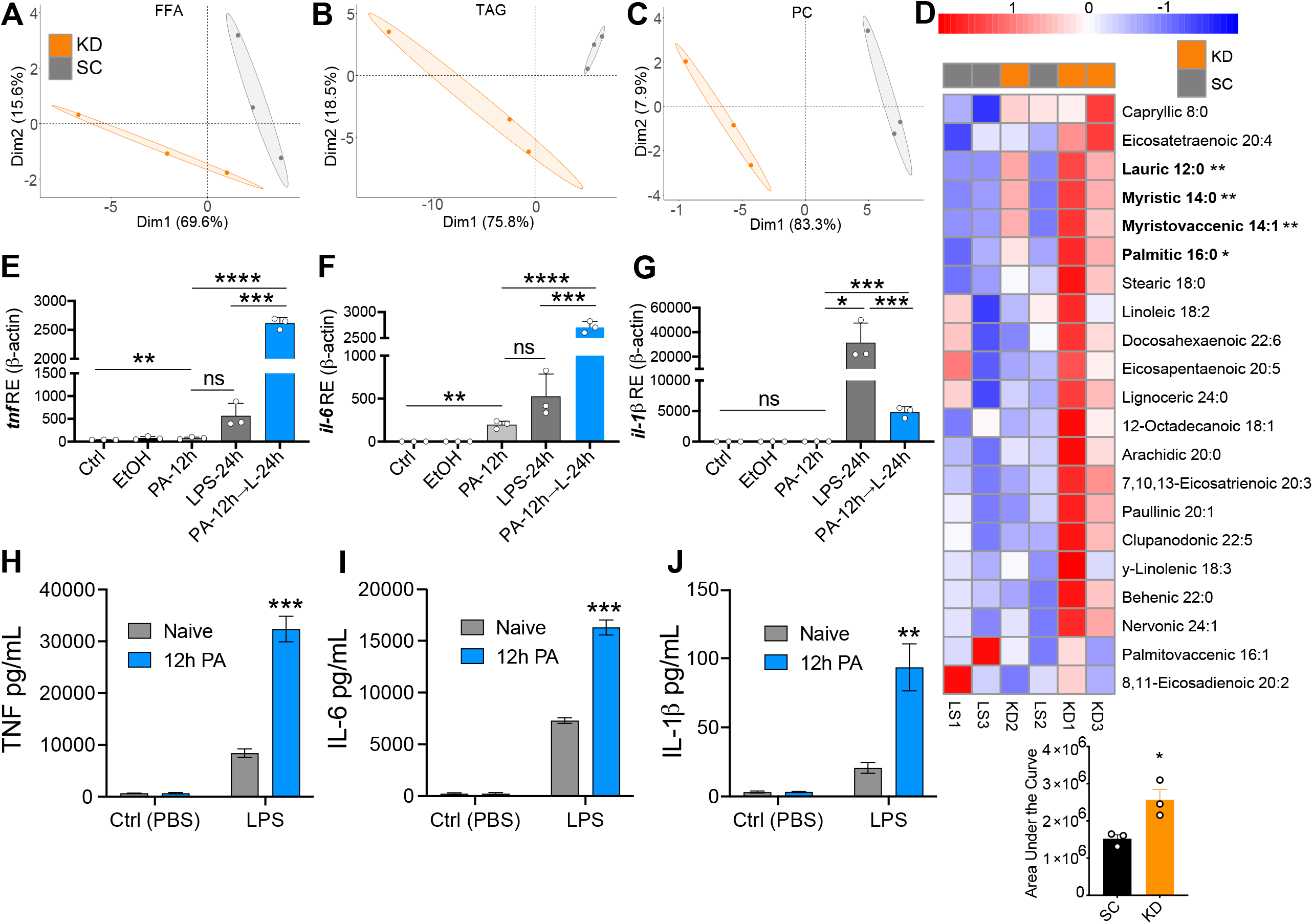
KD alters lipid profiles and PA is mediating a hyper-inflammatory response to secondary challenge with LPS. Data points represent single animal samples and colors represent groups fed SC (black) or KD (orange) diets for two weeks. A 95% confidence ellipse was constructed around the mean point of each group for **(A)** free fatty acids (FFA), **(B)** triglycerides (TAG), and **(C)** phosphatidylcholines (PC). **(D)** Heatmap analysis of free fatty acids in SC and KD mice. Components that are significantly different between the two groups are in bold. Below the heatmap is a comparison of palmitic acid 16:0 peak area detected by LC-MS/MS between SC and KD groups. Statistical significance is determined by unpaired two-tailed t-test between SC and KD groups with n=3 per group. Primary bone marrow-derived macrophages (BMDMs) were isolated from aged-matched female and male mice. BMDMs were plated at 1×10^6^ cells/mL and treated with either ethanol (EtOH; media with 1.69% ethanol), media (Ctrl for LPS), or LPS (10 ng/mL) for 12 h, or palmitic acid (PA stock diluted in 0.83% EtOH; 1 mM PA conjugated to 2% BSA) for 12 h, with and without a secondary challenge with LPS. After indicated time points, RNA was isolated and expression of **(E)** tnf, **(F)** il-6, **(G)** il-1β was measured via qRT-PCR. BMDMs were plated at 1×10^6^ cells/mL and treated with either ethanol (EtOH; media with 1.69% ethanol), media (Naïve), or 1mM PA for 12h followed by PBS (control) or LPS (10ng/mL). Supernatants were assessed via ELISA for **(H)**, TNF, **(I)** IL-6, and **(J)** IL-1β secretion. For all plates, all treatments were performed in triplicate. For all panels, a student’s t-test was used for statistical significance. *, p < 0.05; **, p < 0.01; ***, p < 0.001; ****, p < 0.0001. Error bars show the mean ± SD.

Importantly, we identified a significant increase in multiple circulating FFAs within the KD-fed mice, compared to the SC-fed mice, most of which were SFAs (Fig 3D). Interestingly, in KD-fed mice we found a significant increase in free palmitic acid (PA; C16:0), an immunomodulatory SFA that is found naturally in animal fats, vegetable oils, and human breast milk (*41*), and is enriched in both WD and KD (Fig 3D, Table S1). Additionally, PA-containing SGs, TAGs, and PCs were significantly elevated in KD-fed mice serum, compared to SC-fed mice (Fig S2B-D). These data indicate that KD feeding not only enhances levels of freely circulating PA, but also enhances the frequency PA is incorporated into other lipid species in the blood.

### Palmitic acid enhances macrophage inflammatory response to lipopolysaccharide

Many groups have shown that PA alone induces a modest, but highly reproducible increase in the expression and release of inflammatory cytokines in macrophages and monocytes (*14, 42*). However, it remains unknown if PA can act as a primary inflammatory stimulus to induce a hyperinflammatory response to a secondary heterologous stimulus. Thus, we next wanted to determine if pre-exposure to physiologically relevant concentrations of PA altered the macrophage response to a secondary challenge with LPS. Current literature indicates a wide range of serum PA levels, between 0.7 and 3.6 mM, reflect a high-fat diet in humans (*43-46*). We aimed to use a physiologically relevant concentration of PA reflecting a human host for our *in vitro* studies, thus we treated primary bone marrow-derived macrophages (BMDMs) with and without 1 mM of PA containing 2% bovine serum albumin (BSA) for 12 h, removed the media, subsequently treated with LPS (10 ng/mL) for an additional 24 h, and measured expression and release of TNF, IL-6, and IL-1β. Importantly, the BSA dissolved in the media used for PA treatment solutions was endotoxin- and FA-free to ensure aberrant TLR signaling would not occur via BSA-contamination, and fresh PA was conjugated to BSA-containing media for 30 minutes at 37°C immediately prior to use. We found that BMDMs pre-treated with PA (1 mM) for 12 h expressed significantly higher levels of *tnf* and *il-6* in response to secondary treatment with LPS, compared to naïve BMDMs (Fig 3E, F). *il-1β* expression was significantly lower in cells pre-treated with PA, suggesting a bifurcation in the temporal transcriptional regulation of *tnf/il-6* and *il-1β* by PA (Fig 3G). However, secretion of TNF, IL-6 and IL-1β were all enhanced in BMDMs pre-treated with PA (1 mM) for 12 h and challenged with LPS (Fig 3H-J). We found a similar enhanced *il-6* and *tnf* expression in response to LPS in BMDMs treated with PA (1 mM) for twice the length of exposure (24 h), and *il-1β* expression was again significantly decreased (Fig S4A-C).

Further, we pre-treated BMDMs with a concentration of PA that reflects the lower range of physiologically relevant serum levels and found 0.5 mM of PA induced significantly higher expression of *tnf* and *il-6*, but not *il-1β*, after 12 and 24 h challenge with LPS, compared to naive BMDMs treated with LPS (Fig S4D-I). We conclude that both 12 and 24 h pre-treatments with 0.5 mM or 1 mM of PA conjugated to 2% BSA are sufficient to induce reprogramming of macrophages and alter the response to stimulation with a heterologous ligand. Additionally, these data demonstrate that even serum concentrations of PA that are below the spectrum for humans consuming a high-fat diet pose a risk for inflammatory dysfunction.

Importantly, PA-treatment can induce apoptosis and pyroptosis in various cell types (*47-50*), however we found only 1.9% and 3.4% of cell death after a 12 h or 24 h incubation, respectively, with PA and subsequent 24 h of LPS treatment or control media (Fig S3A, B). These data demonstrate PA pre-treatment of macrophages induces a hyper-inflammatory response to LPS independent of significant cell death, suggesting PA is sensitizing macrophages to secondary inflammatory challenge.

### Diverting ceramide synthesis inhibits the PA-dependent hyper-inflammatory response to LPS

PA treatment diverts cellular metabolism toward the synthesis of the toxic metabolic byproducts: diacylglycerols (DAGs) and ceramide (*51*). PA-induced ceramide has specifically been demonstrated to enhance inflammation (*28, 52-54*). Considering this, we wanted to determine the role of enhanced macrophage ceramide production in driving PA-induced hyper-inflammatory response to LPS. Thus, we treated BMDMs simultaneously with PA (0.5 mM) and a ceramide synthase inhibitor Fumonisin B1 (FB1; 10uM), for 12 h, removed the media, and subsequently treated with LPS (10 ng/mL) for an additional 24 h, and measured release of TNF, IL-6, and IL-1β. We found that BMDMs pre-treated simultaneously with PA (0.5mM) and FB1 (10uM) for 12 h expressed significantly lower levels of TNF, IL-6, and IL-1β secretion in response to LPS, compared to BMDMs pre-treated with only PA (Fig 4A-C). We included a vehicle control for both PA and FB1 [ethanol (EtOH)], and found EtOH did not significantly alter LPS-induced inflammation (Fig 4A-C). We conclude that ceramide synthesis induced by PA is required for the macrophage hyper-inflammatory response to secondary challenge with LPS.

**Fig. 4:**
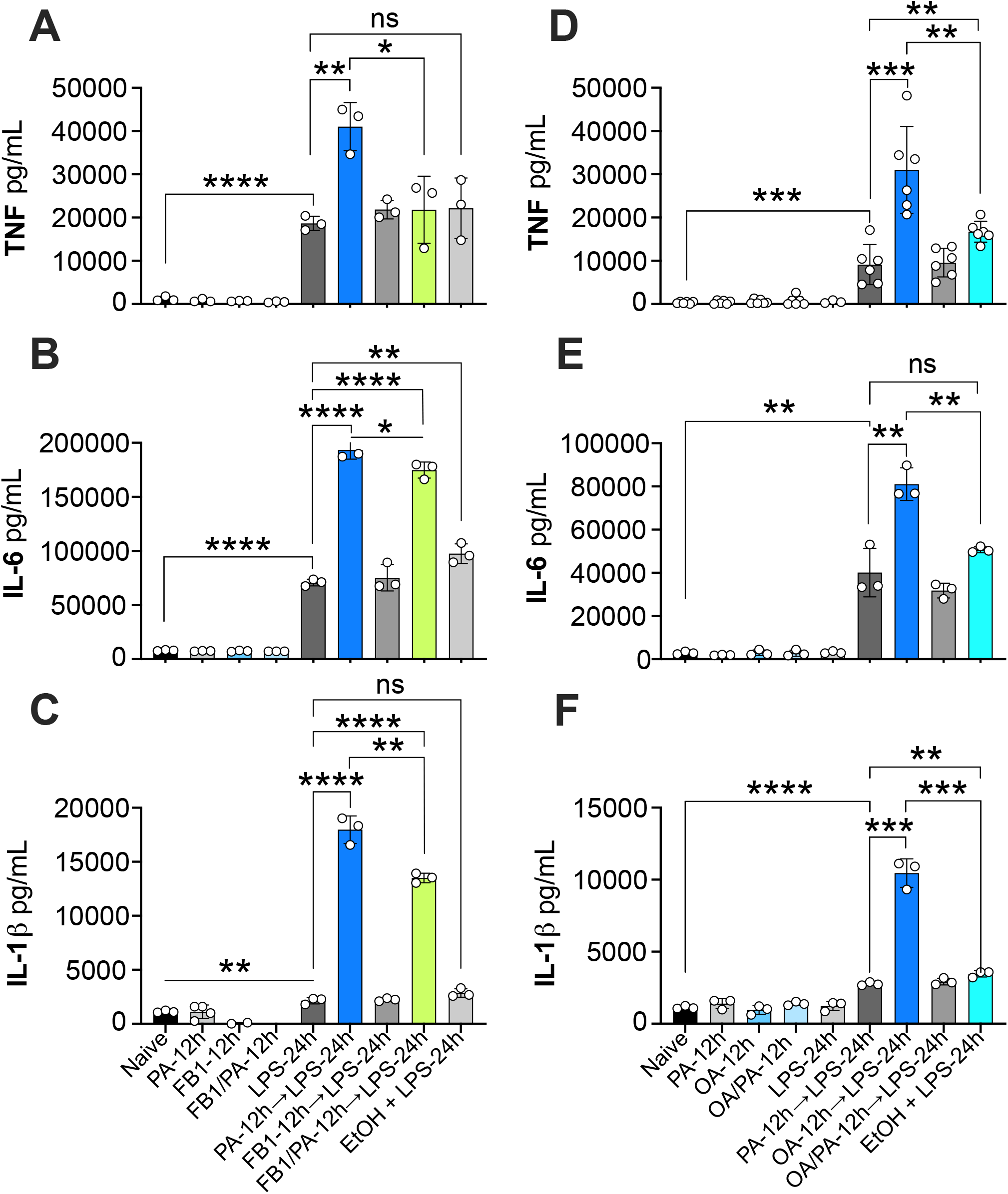
Diverting ceramide synthesis inhibits PA-dependent hyper-inflammatory response to LPS in macrophages. Primary bone marrow-derived macrophages (BMDMs) were isolated from aged-matched female and male mice. BMDMs were plated at 1×10^6^ cells/mL and treated with either media (Ctrl), LPS (10 ng/mL) for 24 h, palmitic acid (PA stock diluted in 0.83% EtOH; 0.5 mM PA conjugated to 2% BSA) for 12 h, Fumonisin B1 (FB1; 10uM; diluted in 0.14% EtOH) alone and simultaneously with PA, oleic acid (OA; 200uM; diluted in endotoxin-free water), or EtOH (0.97% to mimic simultaneous PA/FB1 treatment). Controls for all treatments are shown next to experimental groups treated additionally with LPS (10 ng/mL) for 24 h. Supernatants were assessed via ELISA for **(A), (D)**, TNF, **(B), (E)** IL-6, and **(C), (F)** IL-1β secretion. For all plates, all treatments were performed in triplicate. For all panels, a student’s t-test was used for statistical significance. *, p < 0.05; **, p < 0.01; ***, p < 0.001; ****, p < 0.0001. Error bars show the mean ± SD.

Oleic acid (C18:1) is a mono-unsaturated fatty acid found in animal fats and vegetable oils, and in the presence of PA, diverts lipid metabolism away from ceramide production (*51, 55*). Considering OA and PA are the most prevalent fatty acids found in the human diet and in human serum (*51*), we wanted to test if OA diversion of ceramide synthesis could reverse the PA-dependent hyper-inflammatory response to LPS in macrophages. Thus, we treated BMDMs with OA (200uM), PA (0.5mM), or OA and PA together for 12 h and then with LPS. We found that macrophages simultaneous treated with PA and OA produced significantly lower levels of TNF, IL-6, and IL-1β compared to BMDMs pre-treated with only PA (Fig 4D-F). Together, these data reveal therapeutic and dietary depletion of intracellular ceramides neutralizes the PA-dependent hyper-inflammatory response to LPS in macrophages.

### Palmitic acid is sufficient to increase LPS-induced acute septic shock severity and systemic hyper-inflammation

Considering the drastic effect of PA on macrophage response to secondary challenge with LPS, we next wanted to understand if exposure to PA alone is sufficient to induce a hyper-inflammatory response during LPS-induced acute septic shock. We answered this question by mimicking post-prandial systemic PA levels (1 mM) via a single i.p. injection of ethyl palmitate and then challenging with LPS i.p. (*56*). We found PA-treated mice experienced increased disease severity as indicated by their significant decline in temperature compared to Veh mice (Fig 5A). Similar to WD- and KD-fed mice, PA-treated mice also exhibited 100% mortality, compared to 20% mortality seen in Veh mice (Fig 5B). Importantly, mice injected with PA for shorter time periods (0, 3, and 6 h) and then challenged with LPS did not exhibit increased disease severity or poor survival outcome (Fig S5A, B), concluding that a 12 h pre-treatment with PA is required for an increase in disease severity.

**Fig. 5:**
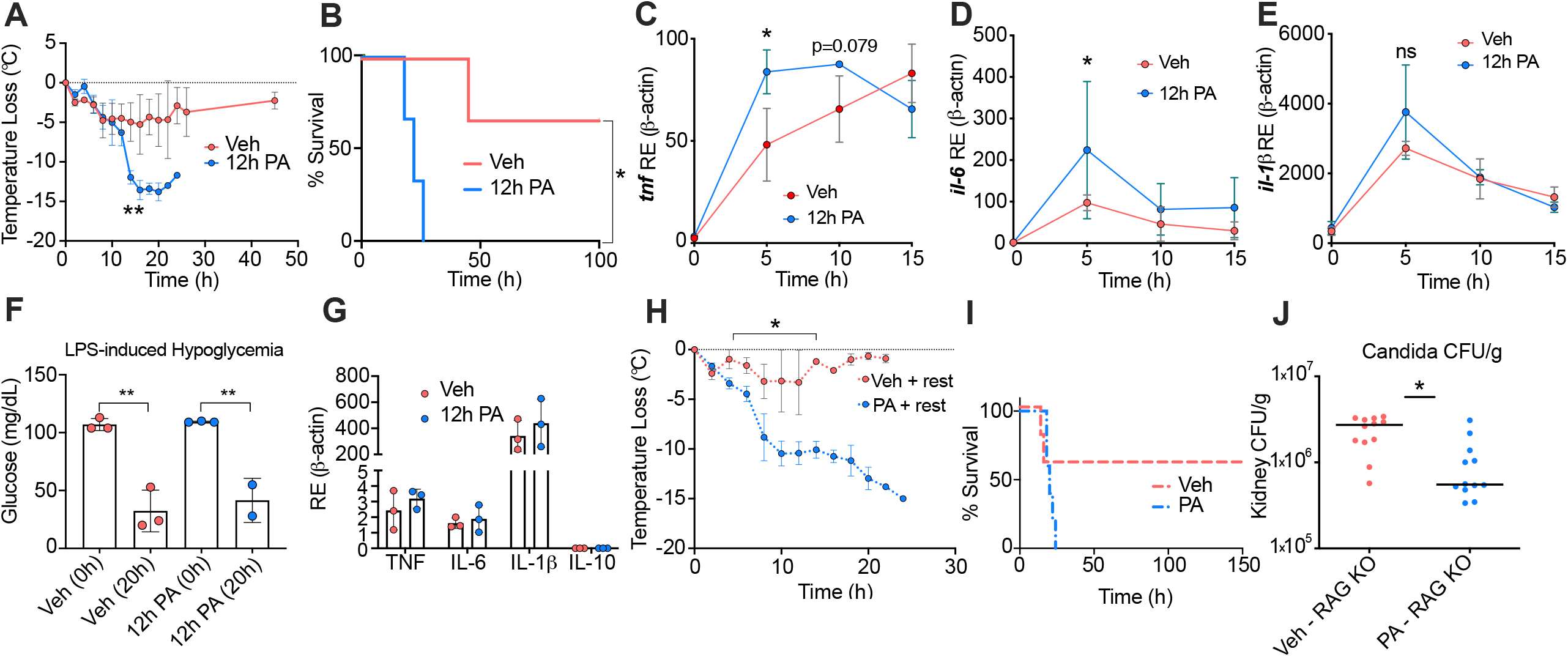
PA is a novel mediator of trained immunity and is sufficient for inducing a hyper-inflammatory response LPS-induced acute septic shock and enhanced clearance of *Candida albicans* infection. Age-matched (4-6 wk) female mice were fed SC for 2 wk and injected i.p. with ethyl palmitate (PA, 750mM) or vehicle (Veh) solutions 12 h before i.p. LPS injections (10 mg/kg). **(A)** Temperature loss was monitored every 2 h as a measure of disease severity or **b** survival. At indicated times 10-20 μL of blood was drawn via the tail vein, RNA was collected, and samples were assessed for expression of **(C)** *tnf*, **(D)** *il-6*, and **(E)** *il-1β* via qRT-PCR. **(F)** At 0 h, before LPS injections, and 20 h p.i. with LPS, blood was collected via the tail vein from Vehicle (Veh) mice, and mice pre-treated with PA 12 h before LPS injections (PA) to measure blood glucose levels. **(G)** Blood was collected via the tail vein from Vehicle (Veh) and PA pre-treated (12 h PA) mice immediately prior to LPS injection and samples were assessed for expression of *tnf, il-6, il-1β*, and *il-10* via qRT-PCR. Additionally, mice were injected i.v. with ethyl palmitate (PA, 750mM) or vehicle (Veh) solutions every day for 9 days and then rested for 7 days before LPS injections (10 mg/kg). **(H)** Temperature loss and **(J)** survival were monitored during endotoxemia. **(J)** Age-matched (8-9 wk) female RAG^-/-^ mice were injected i.v. with ethyl palmitate (PA, 750mM) or vehicle (Veh) solutions 12 h before *C. albicans* infection. Fungal burden of kidneys from Vehicle (Veh) and PA pre-treated (12 h PA) mice 24 h after *C. albicans* infection. For **(A-I)** data are representative of 2 experiments and n=5 mice/group. For **(J)** data are representative of 2 experiments and n=6 mice/group. For **(A), (C-H)**, and **(J)**, a Mann Whitney test was used for pairwise comparisons. For **(B)** and **(I)**, a log-rank Mantel-Cox test was used for survival curve comparison. For all panels, *, *p* < 0.05; **, *p* < 0.01; ***, *p* < 0.001; ****, *p* < 0.0001. Error bars shown mean ± SD.

Next, we measured systemic inflammatory status during disease via the expression of *tnf, il-6, il-1β*, and *il-10* in the blood. We found, similar to WD- and KD-fed mice, the 12 h PA-pre-treated mice showed significantly enhanced expression of *tnf* (15 and 20 h) and *il-6* (5 h) post-LPS challenge, compared to Veh control (Fig 5C, D). Expression of *il-1β* trended upward, but was not significantly up-regulated in 12 h PA-pre-treated mice, compared to Veh-treated mice (Fig 5E). Importantly, as a control we looked at LPS-induced hypoglycemia in PA-treated mice, and 12 h pre-treatment with PA did not alter LPS-induced hypoglycemia (Fig 5F), indicating that diet-dependent hypoglycemic shock was not a driver of acute septic shock severity in PA-treated mice. Thus, exposure to PA to mimic 1 high-fat meal is sufficient to drive enhanced inflammation and disease severity in acutely septic mice, and this effect is dependent on length of PA exposure.

### Enriched dietary PA induces long-lived trained immunity and enhanced clearance of fungal infection

Our data show that PA enhances acute septic shock severity *in vivo*, and enhances inflammatory responses of macrophages to a secondary and heterologous stimulus *in vitro*. This form of regulation resembles non-antigen specific innate immune cell memory; however, it remains unclear if PA is inducing innate immune cell memory by priming or trained immunity. Priming occurs when the first stimulus enhances transcription of inflammatory genes and does not return to basal levels before the secondary stimulation (*57*). In contrast, trained immunity occurs when the first stimulus changes transcription of inflammatory genes, the immune status returns to basal levels, and challenge with a homologous or heterologous stimulus enhances transcription of inflammatory cytokines at much higher levels than those observed during the primary challenge (*57*). Thus, we evaluated the basal level expression of *tnf, il-6*, and *il-1β* in mice treated with 1mM of PA or Veh i.p. for 12h, before stimulation with LPS. Interestingly, we did not see significant differences in *tnf, il-6*, or *il-1β* expression at 12 h of exposure with PA (Fig 5G), which suggests that circulating immune cells of these mice were not in a primed state at these time points prior to LPS injection. These data suggest PA induces trained immunity, and not priming, however the time point of initial inflammation induced by PA remains unknown.

Canonical inducers of trained immunity (e.g., BCG or β-glucan) induce long-lived enhanced innate immune responses to secondary inflammatory stimuli (*58, 59*). Thus, we hypothesized that exposure to a PA bolus would reprogram the inflammatory response *in vivo* and that this program would persist even after mice were “rested” for 1 week. Thus, we injected mice with a vehicle solution or PA (1mM) i.p. once a day for 9 days (to mimic 1 high-fat meal per day) and then rested the mice for 1 wk (Veh or PA→ SC). When challenged with systemic LPS, PA→ SC showed an increase in disease severity and mortality compared to Veh→ SC mice (Fig 5H-I), indicating that PA alone can induce long-lived trained immunity.

Lastly, the most commonly studied models for inducing trained immunity are immunization with BCG or with β-glucan and they have been shown to protect mice from systemic *Candida albicans* infection via lymphocyte-independent epigenetic alterations that lead to decreased kidney fungal burden (*2*). Therefore, we next tested if PA treatment induces lymphocyte-independent clearance of *C. albicans* infection. For these experiments, Rag knockout (Rag^-/-^) mice were treated with a vehicle or PA solution for 12 h and subsequently infected i.v. with 2×10^6^ *C. albicans*. In accordance with canonical trained immunity models, mice treated with PA for 12 h showed a significant decrease in kidney fungal burden compared to Veh mice, 24 h post-infection (Fig 5J). Together, these data find that PA is sufficient to induce long-lived trained immunity *in vivo* that can lead to enhanced lymphocyte-independent host resistance to infection.

## Discussion

In this study we showed that mice fed diets enriched in SFA exhibit hyper-inflammation during LPS-induced acute septic shock and poorer outcomes, compared with mice fed a standard low-SFA diet, independent of the diet-associated microbiome and impact of each diet on LPS-induced hypoglycemia (Fig 1; Fig S1). Strikingly, we found that before LPS treatment, healthy mice fed a diet solely enriched in SFAs (Ketogenic diet; KD) harbored monocytes and splenocytes that were not inherently more inflamed, but when challenged with LPS exhibited increased expression of inflammatory cytokines (Fig 2).

Considering the immunogenic properties of some dietary SFAs enriched in these diets, and that excess dietary SFAs are found circulating throughout the blood and peripheral tissues, we used lipidomics to identify dietary SFAs that may be directly reprogramming innate immune cells to respond more intensely to secondary inflammatory stimuli. Our study identified enriched palmitic acid (C16:0; PA) and PA-associated fatty acids in the blood of KD-fed mice (Fig 3; Fig S2). Thus, we treated macrophages with physiologically relevant concentrations of PA and found that PA induces a hyper-inflammatory response to secondary challenge with LPS (Fig 3; Fig S4). This enhanced response to secondary heterologous stimuli has been shown in previous models of innate immune memory, specifically trained immunity (*4, 60, 61*).

The metabolism of SFAs is a key element of immune system function, and metabolic intermediates enhanced by PA treatment, such as ceramide, serve as signaling lipids in diseases of inflammation (*62*). Mechanistically, we show that inhibiting ceramide synthesis or diverting metabolism away from ceramide synthesis using OA protects macrophages from PA-induced trained immunity, suggesting that dietary intervention may help regulate inflammatory dysregulation during disease. Finally, we conclude that PA-induced trained immunity exacerbates the acute phase of sepsis and correlates with increased mortality, but also enhances resistance to infection independent of mature lymphocytes (Fig 5).

Our findings align with the growing body of evidence indicating that trained immunity is a double-edged sword, where the phenomenon can be beneficial for resistance to infection, but detrimental in the context of inflammatory diseases (*63*). It is known that trained immunity is a key feature of BCG vaccination, which has been shown to enhance resistance to infections, and is a possible mechanism that drives increased resistance to severe COVID-19 in the BCG vaccinated population (*7, 64*). Thus, future research in understanding the plasticity of the PA-regulated trained immune response and enhanced pathogen clearance, and the mechanisms that drive this phenomenon, will be of interest to the larger medical community.

Mechanistically, it is appreciated that PA is not acting as a ligand for the pattern recognition receptor (PRR) TLR4, however the presence of TLR4 (independent of TLR4 signaling capability) is required for PA-dependent inflammation (*14*). Our data and others contribute to the growing evidence that PA is inducing cell intrinsic stress through alterations in metabolism. The crosstalk between glycolytic and oxidative metabolism, and epigenetics, is crucial for trained immunity in human monocytes, and metabolic intermediates of the TCA cycle directly modify histone methylation patterns associated with proinflammatory cytokines upregulated in trained immunity (*4, 65, 66*). While ceramides are known to modify histone acetylation and DNA methylation patterns (*67*), the interplay between ceramide metabolism and epigenetics within innate immune cells has not been explored. Though we have shown that PA-dependent ceramide production leads to innate immune memory, the impact of these alterations on the epigenome remains unknown. Therefore, the influence of ceramide metabolism on epigenetics will be important to consider in future trained immunity studies where PA serves as the primary stimulus.

Interestingly, we find here that immunoparalysis, which is associated with a prolonged septic response and is enhanced in patients with poorer outcomes, is greater in mice fed diets enriched in SFAs (Fig 1) (*32, 33*). However, we found that this SFA-dependent enhanced immunoparalysis is abrogated in germ-free mice, suggesting, for the first time, that the microbial species within the SFA-fed mice may be regulating the late immunoparalytic phase of septic shock. Considering the clinical correlation of immunoparalysis and increased sepsis mortality, it will be imperative to explore the identity of the SFA-dependent microbiome and the host/microbe mechanisms that drive sepsis-associated immunoparalysis.

In conclusion, this unappreciated role of dietary SFAs, specifically PA, may provide insight into the long-lasting immune reprogramming associated with a high-SFA fed population, and lends insight into the complexity of nutritional immunoregulation. Considering the results in this study, we suggest the potential for SFAs such as PA to directly impact innate immune metabolism and epigenetics associated with inflammatory pathways. Thus, our findings are paramount not only for potential dietary interventions, but also treatment of inflammatory diseases exacerbated by metabolic dysfunction in humans.

## Materials and Methods

### Cell lines and reagents

RAW 264.7 macrophages (from ATCC), CASP-1KO BMDMs, BMDMs and BMMs were maintained in DMEM (Gibco) containing L-glutamine, sodium pyruvate, and high glucose supplemented with 10% heat-inactivated fetal bovine serum (FBS; GE Healthcare, SH3039603). BMDMs were also supplemented with 10% macrophage colony-stimulating factor (M-CSF; M-CSF-conditioned media was collected from NIH 3T3 cells expressing M-CSF, generously provided by Denise Monack at Stanford University).

### Generation of BMDMs, BMMs, and splenocytes

Bone marrow-derived macrophages (BMDMs) and bone marrow-derived monocytes (BMMs) were harvested from the femurs and tibias of age-matched (6-8 wk) CO_2_-euthanized female BALB/c mice or male and female C57BL/6J mice. BMDM media was supplemented with 10% macrophage colony-stimulating factor (M-CSF) for differentiation, cells were seeded at 5 × 10^6^ in petri dishes and cultured for 6 days, collected with cold PBS, and frozen in 90% FBS and 10% DMSO in liquid nitrogen for later use. BMMs were isolated from BMDM fraction using EasySep™ Mouse Monocyte Isolation Kit (STEMCELL). Spleens were harvested from age-matched (6-8 wk) CO_2_-euthanized female BALB/c mice, tissue was disrupted using the end of a syringe plunger on a 70 μm cell strainer and rinsed with FACS buffer (PBS + 2mM EDTA). Cells were subjected to red blood cell lysis with RBC lysing buffer (Sigma) followed by neutralization in FACS buffer.

### Treatments

After thawing and culturing for 5 days, BMDMs were pelleted and resuspended in DMEM containing 5% FBS, 2% endotoxin- and fatty acid-free bovine serum albumin (BSA; Proliant Biologicals) and 10% M-CSF. Cells were seeded at 2.5 × 10^5^ cells/well in 24-well tissue-culture plates, treated with EtOH (1.69%, or 0.83%) 10 ng/mL LPS (Ultrapure LPS, *E. coli* 0111:B4, Invivogen), 500 μM or 1 mM palmitic acid (Sigma-Aldrich, PHR1120), 10uM Fumonisin B1 (Sigma-Aldrich, F1147), or 200uM oleic acid (Sigma-Aldrich, O7501). and incubated at 37°C and 5% CO_2_ for 12 or 24 h. Next, cells were treated with an additional 10 ng/mL LPS, and incubated an additional 12 or 24 h. RAW 264.7 macrophages were thawed and cultured for 3-5 days, pelleted and resuspended in DMEM containing 5% FBS and 2% endotoxin- and fatty acid-free BSA, and treated identical to BMDM treatments. BMMs were seeded immediately after harvesting at 4×10^^5^ cells/well in 96-well V-bottom plates in DMEM containing 10% FBS, and treated with LPS for 2 or 24 h. Splenocytes were seeded immediately after harvesting at 1 × 10^5^ cells/well in 96-well V-bottom plates in RPMI media with L-glutamine (Cytiva) containing 10% FBS, and treated with LPS for 2 or 24 h. For all treatments, media was removed and cells were lysed with TRIzol (ThermoFisher), flash-frozen in liquid nitrogen, and stored at -80°C until qRT-PCR analysis. For all plates, all treatments were performed in triplicate.

### Lactate dehydrogenase (LDH) assays

BMDMs were cultured as stated above with culture media, PA, or ethanol in 96-well tissue-culture plates at a concentration of 5 × 10^4^ cells/well and incubated for 12 hours. Cells were treated with PBS or 10 ng/mL LPS in a phenol-red-free Optimem media (ThermoFisher) and incubated an additional 12 or 24 h. Supernatants were collected at the specified time points with LDH release quantified with a CytoTox96 Non-Radioactive Cytotoxicity Assay (Promega). Cytotoxicity was measured per well as a percentage of max LDH release, with background media-only LDH release subtracted. For all plates, all treatments were performed in triplicate.

### Measurement of cell viability

Cell viability was determined by 0.4% Trypan Blue dye exclusion test executed by a TC20 Automated Cell Counter (Bio-Rad).

### Blood RNA extraction and real-time qPCR

Mice were treated with PBS or LPS, and at specified time points 10-20 μL of blood was collected from the tail vein, transferred into 50 μL of RNALater (ThermoFisher Scientific) and frozen at -80°C. RNA extractions were performed using RNeasy Mini Kit (Qiagen) and cDNA was synthesized from RNA samples using SuperScript III First-Strand synthesis system (Invitrogen). Gene specific primers were used to amplify transcripts using SsoAdvanced Universal SYBR Green Supermix (Bio-Rad). A complete list of all primers used, including the names and sequences, is supplied as Supplementary Table 2.

### Enzyme-linked immunosorbent assay (ELISA)

TNF, IL-6, and IL-1β concentrations in mice serum were measured and analyzed using TNF alpha, IL-6, and IL-1β Mouse ELISA Kits (ThermoFisher Scientific), according to the manufacturer’s instructions. Absorbances were measured at a wavelength of 450 nm using a microplate reader (BioTek Synergy HTX). Values below the limit of detection (LOD) of the ELISA were imputed with LOD divided by 2 (LOD/2) values.

### Endotoxin-induced model of sepsis

Age-matched (6-8 wk) female BALB/c mice were anesthetized with isoflurane and injected subcutaneously with ID transponders (Bio Medic Data Systems). 2 wk post diet change, and 1 wk post ID transponder injection, mice were stimulated with a single injection of 6-10 mg/kg LPS reconstituted in endotoxin-free LAL reagent water (Invivogen) and diluted in PBS for a total volume of 200 μL. Control mice received corresponding volumes of PBS. Progression of disease was monitored every 2 h after LPS injection for clinical signs of endotoxin shock based on weight, coat and eyes appearance, level of consciousness and locomotor activity. Temperature was recorded using a DAS-8007 thermo-transponder wand (Bio Medic Data Systems). For PA injections, a solution of 750 mM ethyl palmitate (Millipore Sigma), 1.6% lecithin (Sigma-Aldrich) and 3.3% glycerol was made in endotoxin-free LAL reagent water (Lonza). The lecithin-glycerol-water solution was used as a vehicle, and mice were injected with 200 μL of the vehicle as a control, or ethyl palmitate solution to increase serum PA levels to 1 mM (*56*).

### Mouse diets, glucose, and ketones

Six-week-old female mice were fed soft, irradiated chow and allowed to acclimate to research facility undisturbed for one week. Chow was replaced by Western Diet (Envigo, TD.88137), Ketogenic Diet (Envigo, TD.180423), or Standard Chow (Envigo, TD.08485) and mice were fed *ad libitum* for two weeks before induction of sepsis. For Ketogenic Diet, food was changed daily. For Western Diet, food was changed every 72 hours. Ketones and blood glucose were measured weekly and immediately prior to LPS injections with blood collected from the tail vein using Blood Ketone & Glucose Testing Meter (Keto-Mojo), or with urine collected on ketone indicator strips (One Earth Health, Ketone Test Strips).

### Statistics analysis

Mann Whitney, Mantel-Cox, and student’s t-tests were carried out with GraphPad Prism 9.0 software.

### Ethical approval of animal studies

All animal studies were performed in accordance with National Institutes of Health (NIH) guidelines, the Animal Welfare Act, and US federal law. All animal experiments were approved by the Oregon Health and Sciences University (OHSU) Department of Comparative Medicine or Oregon State University (OSU) Animal Program Office and were overseen by the Institutional Care and Use Committee (IACUC) under Protocol IDs #IP00002661 & IP00001903 at OHSU and #5091 at OSU. Conventional animals were housed in a centralized research animal facility certified by OHSU. Conventional 6-10 wk-aged female BALB/c mice (Jackson Laboratory 000651) were used for the sepsis model, and isolation of BMDMs, BMMs, and splenocytes. GF male and female C57BL/6 mice (Oregon State University; bred in house) between 14 and 23 wk old were used for the GF sepsis model. BALB/c Rag1^-/-^ mice between 8 and 24 wk were infected i.v. with 2×10^6^ CFUs of *C. albicans* SC5314 (ATCC #MYA-2876) and kidney fungal burden was assessed 24 h post-infection. Kidneys were harvested 24 h post-infection and homogenized organs were plated in serial dilutions on Yeast Peptone Dextrose plates to assess fungal burden.

### Lipidomics PCA Analysis

Mice on specialized diets were sacrificed at the indicated time points after PBS or LPS treatment with 300-600µL of blood collected via cardiac puncture into heparinized tubes. Blood samples were centrifuged for 15 minutes at 2,500rpm at 4°C and plasma was transferred to a new tube before storage at -80°C. Plasma samples were analyzed via LC-MS/MS. Lipidomic data sets were scaled using the *scale* function and principal component analyses were performed using the *prcomp* function from the stats package in R Version 3.6.2. Visualization of PCAs and biplots was performed with the *fviz_pca_ind* and *fviz_pca_biplot* functions from the factoextra package and with the *ggplot2* package (*68, 69*). For each diet group, 95% confidence ellipses were plotted around the group mean using the *coord*.*ellipse* function from the FactoMineR package (*70*). Heatmaps were created using the *pheatmap* package (*71*).

## Acknowledgements

We would like to thank Ajesh Saini, a student in the Napier Lab, for his contributions in carrying out the ELISA data within this manuscript. This study was supported by National Institute of General Medical Sciences (NIGMS) grant 5R35GM133804-02 to B.A.N.

---

**Supplementary Figure 1.**
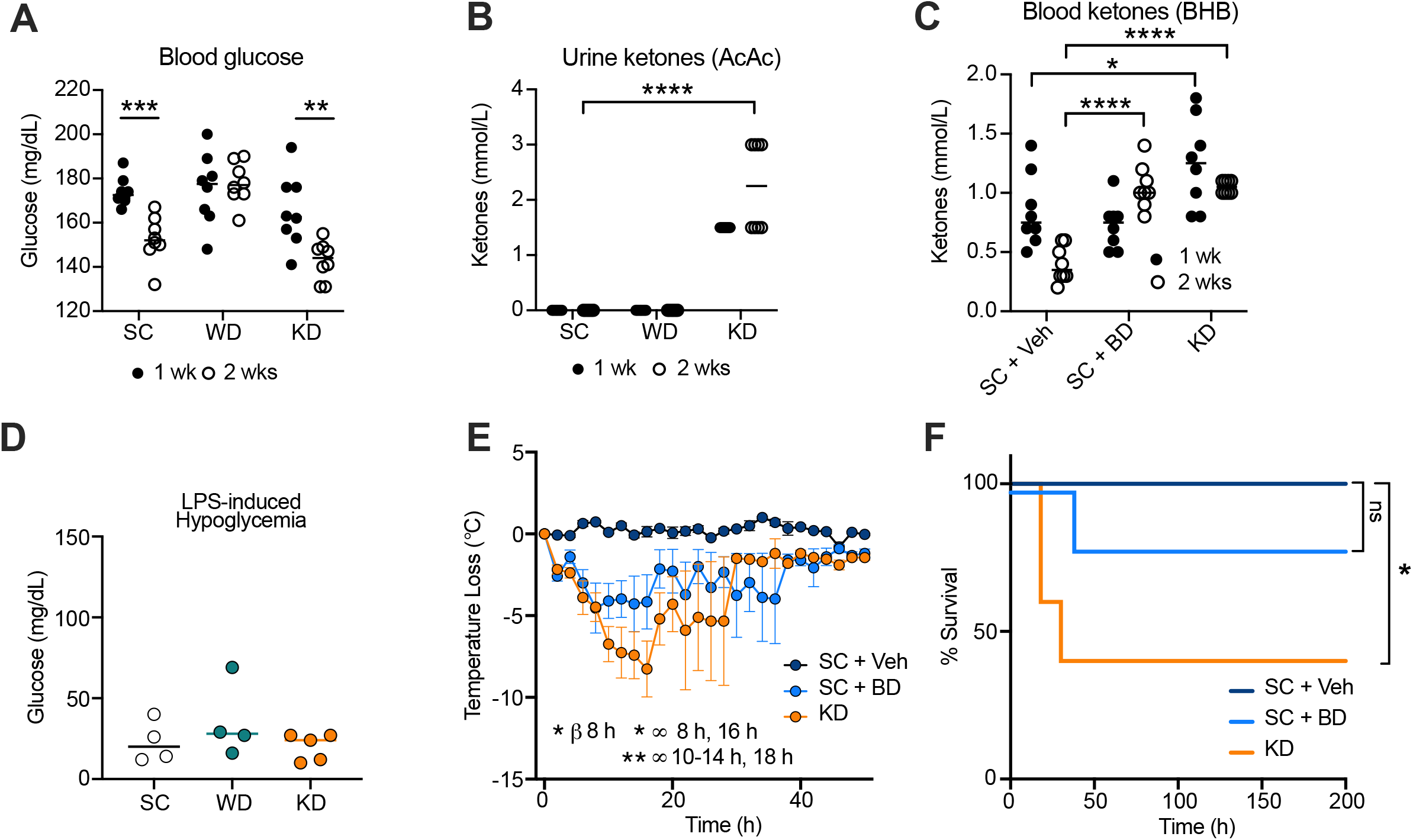
Increase in disease severity in KD mice is independent of ketosis. Age-matched mice were fed a SC, WD, or KD for 2 wk. At 1 wk and 2 wk **a** blood was collected via the tail vein to measure blood glucose levels using a glucose testing meter (Keto-Mojo) and **b** urine was collected on ketone indicator strips to measure levels of systemic acetoacetate (AcAc). **c** Age-matched mice were fed a SC supplemented with 1,3-butanediol (SC + BD) or with the saccharine vehicle solution as a control (SC + Veh), or KD for 2 wk. At 1 wk and 2 wk blood was collected via the tail vein to measure levels of systemic *β*-hydroxybutyrate (BHB) using a ketone testing meter (Keto-Mojo). **d** At 2 wk, mice were injected i.p. with LPS and 25 h p.i. blood glucose levels were measured as stated in **a. e** Temperature loss and **f** survival were monitored every 2 h. **a-c** All experiments were run 8 times and data are representative of 1 experiment. *n* = 5 mice/group. A Mann-Whitney U test was used for pairwise comparisons. **d, e** All experiments were run 3 times and data are representative of 1 experiment. A student’s t-test was used for statistical significance. For **e**, β symbols indicate SC + Veh vs. SC + BD significance and ∞ symbols indicate SC + Veh vs. KD significance. For **f**, a log-rank Mantel-Cox test was used for survival curve comparison. For all panels, a *p-*value of less than 0.05 was considered significant (*, *p* < 0.05; **, *p* < 0.01; ***, *p* < 0.001; ****, *p* < 0.0001).

**Supplementary Figure 2.**
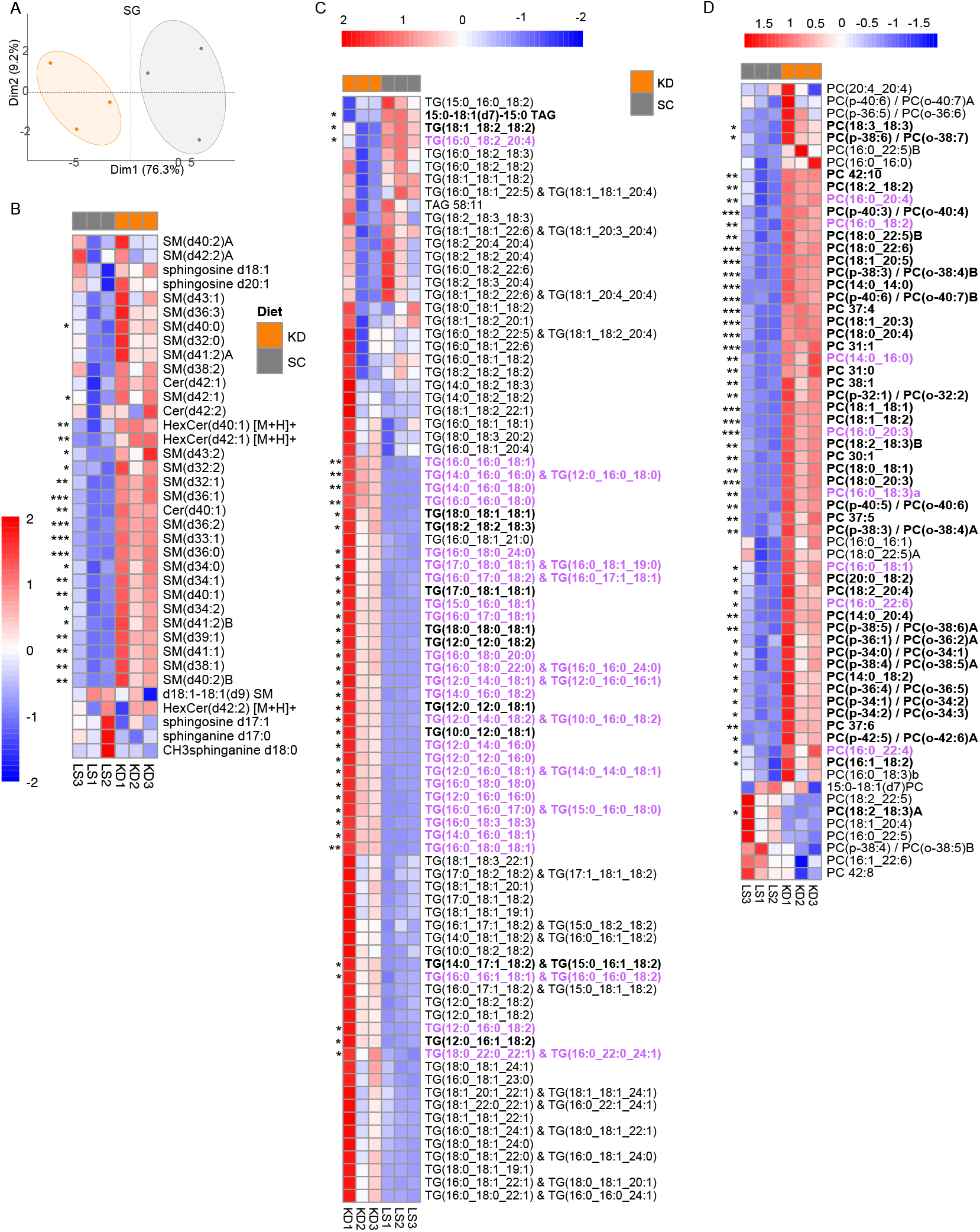
Principal component analysis (PCA) and heatmap analysis of sphingolipid lipidomic data in mouse plasma samples. **a** Data points represent single animal samples and colors represent groups fed SC (black) or KD (orange) diets for two weeks and a 95% confidence ellipse was constructed around the mean point of each group. Heatmap analysis of **b** sphingolipids (SM), **c** triglycerides (TG), and **d** phosphatidylcholines (PC) in SC and KD groups. Lipid components containing 16:0 palmitic chains are highlighted in purple and components that are significantly different between the two groups are in bold. Statistical significance determined by unpaired two-tailed t-test between SC and KD groups. *, P < 0.05; **, P < 0.01; ***, P < 0.001. n=3 per group.

**Supplementary Figure 3.**
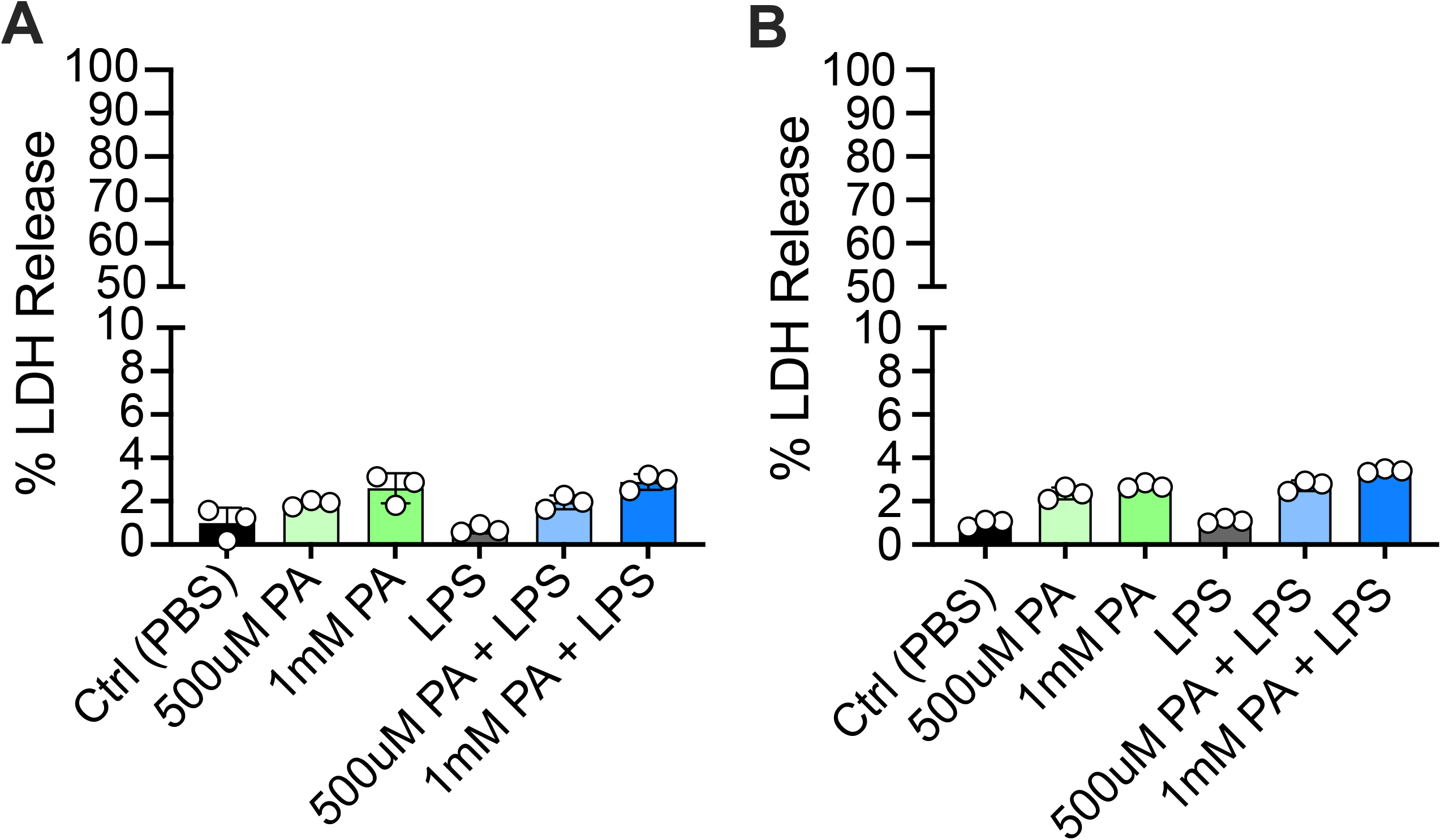
Cytotoxicity in PA-treated BMDMs. Black6 BMDMs were cultured and plated in 96-well plates and treated with EtOH (1.69%) as a carrier control, palmitic acid (PA; 1mM), or BMDM Media for 12h and then incubated with PBS (control), BMDM Media, or LPS (10ng/mL) for **a** 12 h or **b** 24h and release of lactate dehydrogenase (LDH) was measured via CytoTox96 Non-Radioactive Cytotoxicity Assay (Promega). Cytotoxicity was assessed per well as a percentage of max LDH release, with background absorption subtracted. For all plates, all treatments were performed in triplicate. For all panels, data was normalized to control treatments, and a student’s t-test was used for statistical significance. *, p < 0.05; **, p < 0.01; ***, p< 0.001; ****, p < 0.0001. For all panels, error bars show the mean ± SD.

**Supplementary Figure 4.**
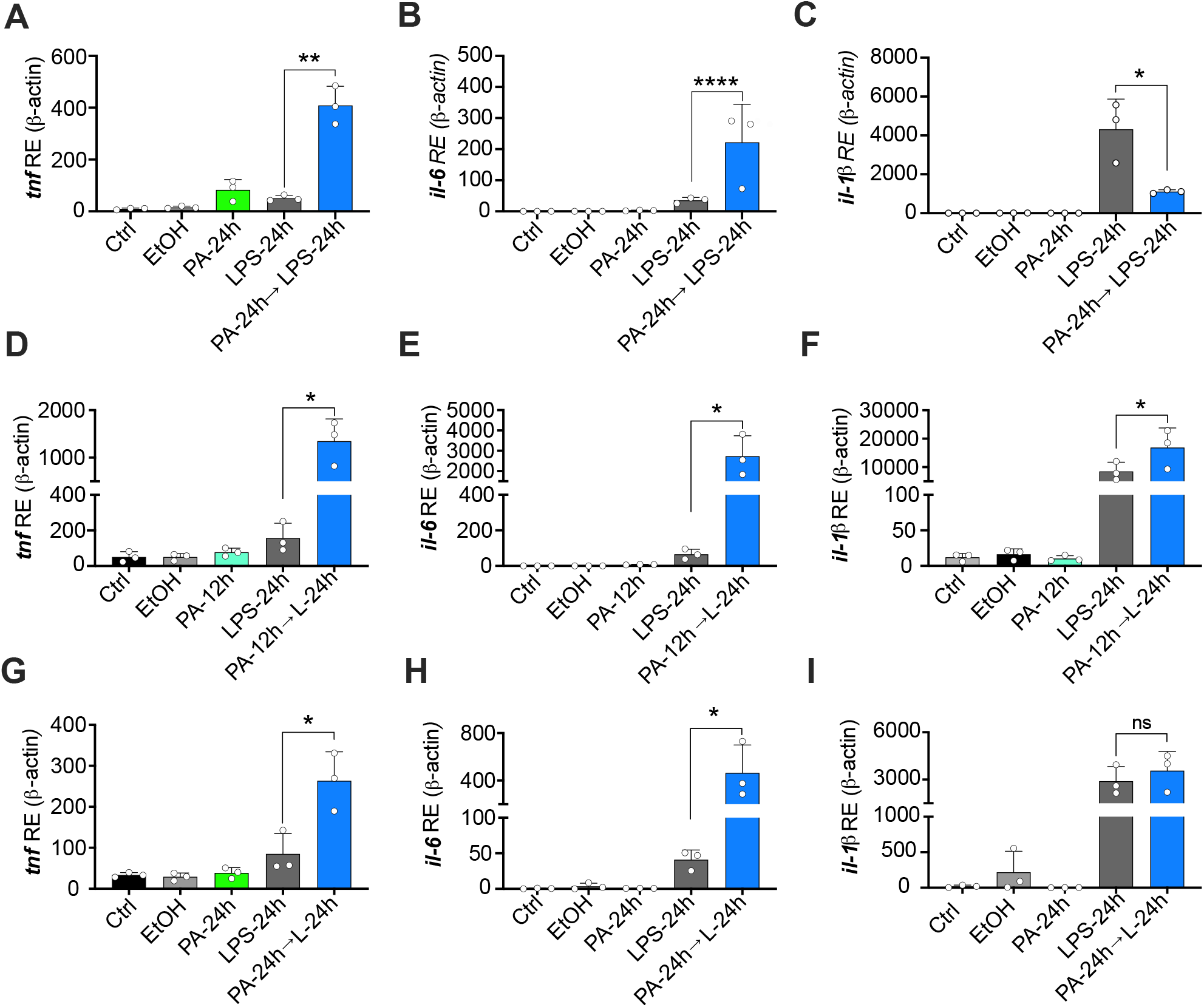
Physiological levels of Palmitic Acid (PA) induce a hyper-inflammatory response to secondary challenge with LPS in macrophages. Primary bone marrow-derived macrophages (BMDMs) were isolated from aged-matched female and male mice. BMDMs were plated at 1×10^6^ cells/mL and treated with either ethanol (EtOH; media with 1.69% ethanol), media (Ctrl for LPS), LPS (10 ng/mL) for 12 or 24 h, or palmitic acid (PA; 0.5mM; diluted in 1.69% EtOH) for 12 h, with and without a secondary challenge with LPS. Next, BMDMs were treated with either EtOH, media, LPS (10 ng/mL) for 24 h, **a-f** PA (1mM; diluted in 1.69% EtOH) for 12 h with and without a secondary challenge with LPS. After indicated time points, RNA was isolated and expression of **a, d** *tnf*, **b, e** *il-6*, and **c, f** *il-1β* was measured via qRT-PCR. Caspase-1 KO BMDMs were plated at 1×10^6^ cells/mL and treated as described above. After indicated time points, RNA was isolated and expression of **g, j** *tnf*, **h, k** *il-6*, and **i, l** *il-1β* was measured via qRT-PCR. Supernatants were assessed via ELISA for **(H)**, TNF, **(I)** IL-6, and **(J)** IL-1β secretion. For all plates, all treatments were performed in triplicate. For all panels, a student’s t -test was used for statistical significance. *, p < 0.05; **, p < 0.01; ***, p< 0.001; ****, p < 0.0001. For all panels, error bars show the mean ± SD.

**Supplementary Figure 5.**
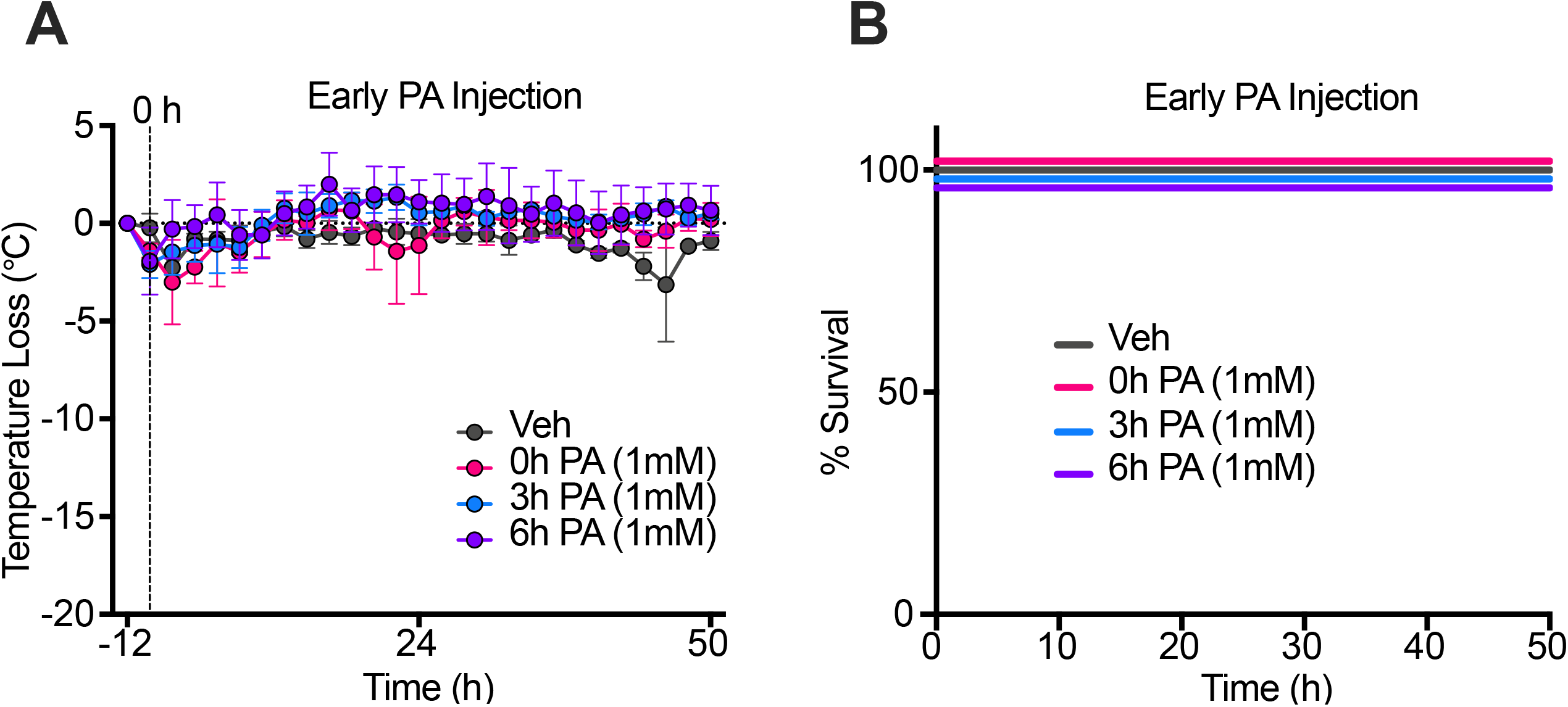
Palmitic acid-induced trained immunity in endotoxemia is time-dependent. Age-matched (4-6 wk) mice were fed SC for 2 wk and injected i.p. with ethyl palmitate (PA, 750mM in 1.6% lecithin and 3.3% glycerol in endotoxin-free LAL reagent water) or vehicle (Veh; 1.6% lecithin and 3.3% glycerol in endotoxin-free LAL reagent water). At 0, 3, and 6 h after PA injection, endotoxemia was induced vi a single i.p. injection of LPS (10 mg/kg). **a** Temperature loss and **b** survival were monitored every 2 h. Data are representative of 1 experiment, *n* = 3 mice/group.

## Supplemental Tables

**Supplemental Table 1.** Diet compositions (values represent percentage of total kcal).

| Diet | Envigo Cat. No. | Fat | Carb | Protein | kcal/g |
| --- | --- | --- | --- | --- | --- |
| Standard Chow | TD.08485 | 13 (3% PA) | 61.3 | 19.1 | 3.6 |
| Western Diet | TD.88137 | 42 (12% PA) | 42.7 | 15.2 | 4.5 |
| Ketogenic Diet | TD.180423 | 90.5 (23% PA) | 0.4 | 9.1 | 6.8 |

**Supplemental Table 2.**
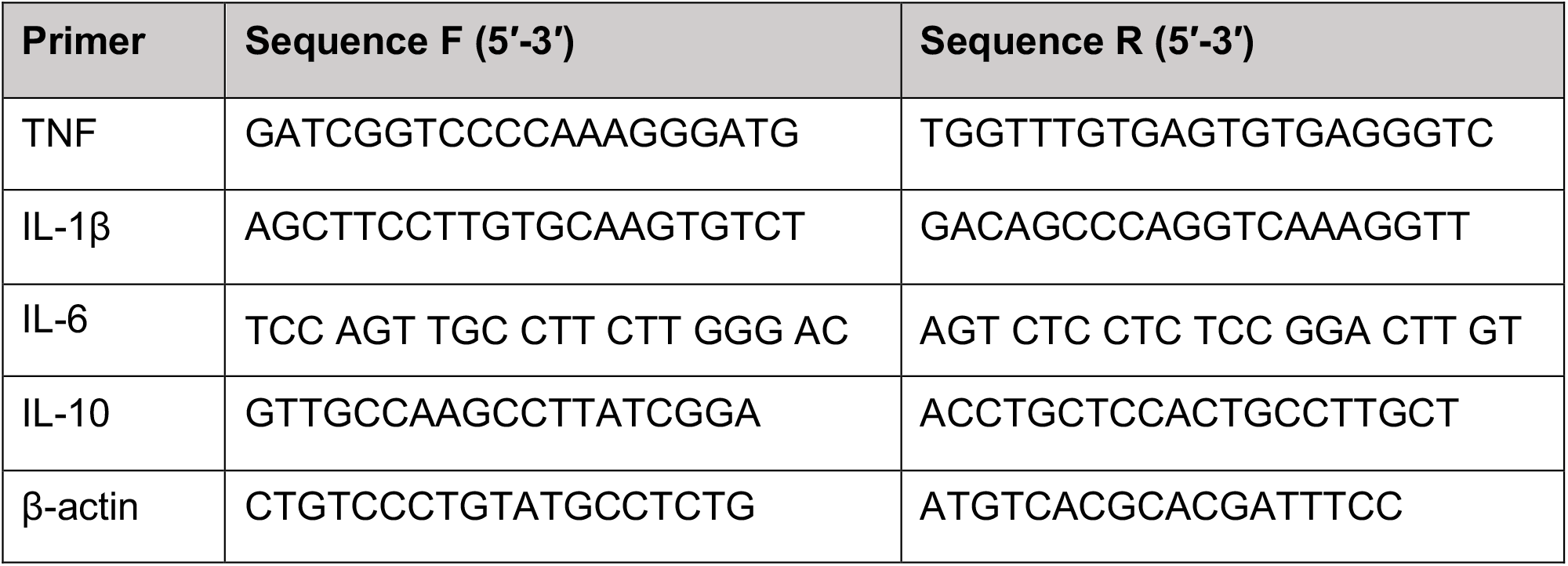
List of primers used in this study.

## References

1. M. G. Netea et al., Defining trained immunity and its role in health and disease. Nat Rev Immunol 20, 375–388 (2020).

2. J. Kleinnijenhuis et al., Bacille Calmette-Guerin induces NOD2-dependent nonspecific protection from reinfection via epigenetic reprogramming of monocytes. Proc Natl Acad Sci U S A 109, 17537–17542 (2012).

3. J. Quintin et al., Candida albicans infection affords protection against reinfection via functional reprogramming of monocytes. Cell Host Microbe 12, 223–232 (2012).

4. S. Saeed et al., Epigenetic programming of monocyte-to-macrophage differentiation and trained innate immunity. Science 345, 1251086 (2014).

5. H. Lanz-Mendoza, J. Contreras-Garduno, Innate immune memory in invertebrates: Concept and potential mechanisms. Dev Comp Immunol 127, 104285 (2022).

6. N. Katzmarski et al., Transmission of trained immunity and heterologous resistance to infections across generations. Nat Immunol 22, 1382–1390 (2021).

7. M. G. Netea et al., Trained immunity: A program of innate immune memory in health and disease. Science 352, aaf1098 (2016).

8. C. Covian, A. Retamal-Diaz, S. M. Bueno, A. M. Kalergis, Could BCG Vaccination Induce Protective Trained Immunity for SARS-CoV-2? Front Immunol 11, 970 (2020).

9. D. K. Meena et al., Beta-glucan: an ideal immunostimulant in aquaculture (a review). Fish Physiol Biochem 39, 431–457 (2013).

10. M. van Splunter et al., Induction of Trained Innate Immunity in Human Monocytes by Bovine Milk and Milk-Derived Immunoglobulin G. Nutrients 10, (2018).

11. B. A. Swinburn et al., The global obesity pandemic: shaped by global drivers and local environments. Lancet 378, 804–814 (2011).

12. B. M. Popkin, Global nutrition dynamics: the world is shifting rapidly toward a diet linked with noncommunicable diseases. Am J Clin Nutr 84, 289–298 (2006).

13. A. Christ, E. Latz, The Western lifestyle has lasting effects on metaflammation. Nat Rev Immunol 19, 267–268 (2019).

14. G. I. Lancaster et al., Evidence that TLR4 Is Not a Receptor for Saturated Fatty Acids but Mediates Lipid-Induced Inflammation by Reprogramming Macrophage Metabolism. Cell Metab 27, 1096–1110.e1095 (2018).

15. C. N. Lumeng, J. L. Bodzin, A. R. Saltiel, Obesity induces a phenotypic switch in adipose tissue macrophage polarization. J Clin Invest 117, 175–184 (2007).

16. P. J. Meikle, S. A. Summers, Sphingolipids and phospholipids in insulin resistance and related metabolic disorders. Nat Rev Endocrinol 13, 79–91 (2017).

17. A. W. B. Reyes et al., Protection of palmitic acid treatment in RAW264.7 cells and BALB/c mice during Brucella abortus 544 infection. Journal of Veterinary Science 22, (2021).

18. B. A. Napier et al., Western diet regulates immune status and the response to LPS-driven sepsis independent of diet-associated microbiome. Proc Natl Acad Sci U S A 116, 3688–3694 (2019).

19. E. L. Goldberg et al., Ketogenic diet activates protective γδ T cell responses against influenza virus infection. Sci Immunol 4, (2019).

20. B. A. Napier et al., Complement pathway amplifies caspase-11-dependent cell death and endotoxin-induced sepsis severity. J Exp Med, (2016).

21. H. Saito, E. R. Sherwood, T. K. Varma, B. M. Evers, Effects of aging on mortality, hypothermia, and cytokine induction in mice with endotoxemia or sepsis. Mech Ageing Dev 124, 1047–1058 (2003).

22. C. F. Raetzsch et al., Lipopolysaccharide inhibition of glucose production through the Toll-like receptor-4, myeloid differentiation factor 88, and nuclear factor kappa b pathway. Hepatology 50, 592–600 (2009).

23. J. P. Filkins, R. P. Cornell, Depression of hepatic gluconeogenesis and the hypoglycemia of endotoxin shock. Am J Physiol 227, 778–781 (1974).

24. A. J. Lewis, C. W. Seymour, M. R. Rosengart, Current Murine Models of Sepsis. Surg Infect (Larchmt) 17, 385–393 (2016).

25. A. Wang et al., Opposing Effects of Fasting Metabolism on Tissue Tolerance in Bacterial and Viral Inflammation. Cell 166, 1512–1525.e1512 (2016).

26. Y. V. Radzyukevich, N. I. Kosyakova, I. R. Prokhorenko, Participation of Monocyte Subpopulations in Progression of Experimental Endotoxemia (EE) and Systemic Inflammation. Journal of Immunology Research 2021, 1–9 (2021).

27. L. Zhang et al., Splenocyte Apoptosis and Autophagy Is Mediated by Interferon Regulatory Factor 1 During Murine Endotoxemia. Shock 37, 511–517 (2012).

28. E. A. Schwartz et al., Nutrient modification of the innate immune response: a novel mechanism by which saturated fatty acids greatly amplify monocyte inflammation. Arterioscler Thromb Vasc Biol 30, 802–808 (2010).

29. C. Fang et al., Differential regulation of lipopolysaccharide-induced IL-1beta and TNF-alpha production in macrophages by palmitate via modulating TLR4 downstream signaling. Int Immunopharmacol 103, 108456 (2021).

30. S. C. Cheng et al., Broad defects in the energy metabolism of leukocytes underlie immunoparalysis in sepsis. Nat Immunol 17, 406–413 (2016).

31. C. Nedeva, J. Menassa, H. Puthalakath, Sepsis: Inflammation Is a Necessary Evil. Front Cell Dev Biol 7, 108 (2019).

32. C. A. Gogos, E. Drosou, H. P. Bassaris, A. Skoutelis, Pro-versus anti-inflammatory cytokine profile in patients with severe sepsis: a marker for prognosis and future therapeutic options. J Infect Dis 181, 176–180 (2000).

33. J. T. van Dissel, P. van Langevelde, R. G. Westendorp, K. Kwappenberg, M. Frölich, Anti-inflammatory cytokine profile and mortality in febrile patients. Lancet 351, 950–953 (1998).

34. R. M. Dougherty, C. Galli, A. Ferro-Luzzi, J. M. Iacono, Lipid and phospholipid fatty acid composition of plasma, red blood cells, and platelets and how they are affected by dietary lipids: a study of normal subjects from Italy, Finland, and the USA. Am J Clin Nutr 45, 443–455 (1987).

35. C. M. Skeaff, L. Hodson, J. E. McKenzie, Dietary-induced changes in fatty acid composition of human plasma, platelet, and erythrocyte lipids follow a similar time course. J Nutr 136, 565–569 (2006).

36. N. Zollner, F. Tato, Fatty acid composition of the diet: impact on serum lipids and atherosclerosis. Clin Investig 70, 968–1009 (1992).

37. J. Choi et al., Comprehensive analysis of phospholipids in the brain, heart, kidney, and liver: brain phospholipids are least enriched with polyunsaturated fatty acids. Mol Cell Biochem 442, 187–201 (2018).

38. I. S. S. A, A. B. C, J. S. A, Changes in Plasma Free Fatty Acids Associated with Type-2 Diabetes. Nutrients 11, (2019).

39. E. Sokolowska, A. Blachnio-Zabielska, The Role of Ceramides in Insulin Resistance. Front Endocrinol (Lausanne) 10, 577 (2019).

40. M. Papadimitriou-Olivgeris et al., The Role of Obesity in Sepsis Outcome among Critically Ill Patients: A Retrospective Cohort Analysis. Biomed Res Int 2016, 5941279 (2016).

41. A. Mancini et al., Biological and Nutritional Properties of Palm Oil and Palmitic Acid: Effects on Health. Molecules 20, 17339–17361 (2015).

42. J. Korbecki, K. Bajdak-Rusinek, The effect of palmitic acid on inflammatory response in macrophages: an overview of molecular mechanisms. Inflamm Res 68, 915–932 (2019).

43. L. Liu et al., Targeted metabolomic analysis reveals the association between the postprandial change in palmitic acid, branched-chain amino acids and insulin resistance in young obese subjects. Diabetes Res Clin Pract 108, 84–93 (2015).

44. S. F. Gallego, M. Hermansson, G. Liebisch, L. Hodson, C. S. Ejsing, Total Fatty Acid Analysis of Human Blood Samples in One Minute by High-Resolution Mass Spectrometry. Biomolecules 9, (2018).

45. C. D. C. Buchanan et al., Analysis of major fatty acids from matched plasma and serum samples reveals highly comparable absolute and relative levels. Prostaglandins, Leukotrienes and Essential Fatty Acids 168, 102268 (2021).

46. M. Perreault et al., A distinct fatty acid profile underlies the reduced inflammatory state of metabolically healthy obese individuals. PLoS One 9, e88539 (2014).

47. N. M. Borradaile et al., Disruption of endoplasmic reticulum structure and integrity in lipotoxic cell death. J Lipid Res 47, 2726–2737 (2006).

48. H. Li et al., Adipocyte Fatty Acid-Binding Protein Promotes Palmitate-Induced Mitochondrial Dysfunction and Apoptosis in Macrophages. Front Immunol 9, 81 (2018).

49. L. D. Ly et al., Oxidative stress and calcium dysregulation by palmitate in type 2 diabetes. Exp Mol Med 49, e291 (2017).

50. L. Tao et al., RIP1 kinase activity promotes steatohepatitis through mediating cell death and inflammation in macrophages. Cell Death Differ 28, 1418–1433 (2021).

51. X. Palomer, J. Pizarro-Delgado, E. Barroso, M. Vázquez-Carrera, Palmitic and Oleic Acid: The Yin and Yang of Fatty Acids in Type 2 Diabetes Mellitus. Trends in Endocrinology & Metabolism 29, 178–190 (2018).

52. J. Jin et al., Acid sphingomyelinase plays a key role in palmitic acid-amplified inflammatory signaling triggered by lipopolysaccharide at low concentrations in macrophages. Am J Physiol Endocrinol Metab 305, E853–867 (2013).

53. J. D. Schilling et al., Palmitate and Lipopolysaccharide Trigger Synergistic Ceramide Production in Primary Macrophages. Journal of Biological Chemistry 288, 2923–2932 (2013).

54. Y. Zhang et al., Adipose Fatty Acid Binding Protein Promotes Saturated Fatty Acid-Induced Macrophage Cell Death through Enhancing Ceramide Production. J Immunol 198, 798–807 (2017).

55. L. L. Listenberger et al., Triglyceride accumulation protects against fatty acid-induced lipotoxicity. Proc Natl Acad Sci U S A 100, 3077–3082 (2003).

56. K. Eguchi et al., Saturated fatty acid and TLR signaling link beta cell dysfunction and islet inflammation. Cell Metab 15, 518–533 (2012).

57. M. Divangahi et al., Trained immunity, tolerance, priming and differentiation: distinct immunological processes. Nat Immunol 22, 2–6 (2021).

58. E. Kaufmann et al., BCG Educates Hematopoietic Stem Cells to Generate Protective Innate Immunity against Tuberculosis. Cell 172, 176–190 e119 (2018).

59. M. G. Netea et al., Defining trained immunity and its role in health and disease. Nat Rev Immunol 20, 375–388 (2020).

60. S. Bekkering et al., Oxidized low-density lipoprotein induces long-term proinflammatory cytokine production and foam cell formation via epigenetic reprogramming of monocytes. Arterioscler Thromb Vasc Biol 34, 1731–1738 (2014).

61. A. Christ et al., Western Diet Triggers NLRP3-Dependent Innate Immune Reprogramming. Cell 172, 162–175.e114 (2018).

62. S. Galadari, A. Rahman, S. Pallichankandy, A. Galadari, F. Thayyullathil, Role of ceramide in diabetes mellitus: evidence and mechanisms. Lipids Health Dis 12, 98 (2013).

63. A. R. DiNardo, M. G. Netea, D. M. Musher, Postinfectious Epigenetic Immune Modifications - A Double-Edged Sword. N Engl J Med 384, 261–270 (2021).

64. L. E. Escobar, A. Molina-Cruz, C. Barillas-Mury, BCG vaccine protection from severe coronavirus disease 2019 (COVID-19). Proc Natl Acad Sci U S A 117, 17720–17726 (2020).

65. R. J. Arts et al., Glutaminolysis and Fumarate Accumulation Integrate Immunometabolic and Epigenetic Programs in Trained Immunity. Cell Metab 24, 807–819 (2016).

66. D. G. Ryan, L. A. J. O’Neill, Krebs Cycle Reborn in Macrophage Immunometabolism. Annu Rev Immunol 38, 289–313 (2020).

67. G. D. Silva, F. C. Brochers-Lacchini, A. M. Leopoldino, How do sphingolipids play a role in epigenetic mechanisms and gene expression? Epigenomics, (2021).

68. A. K. F. Mundt et al. (2017).

69. Wickham et al. (Springer International Publishing, 2016).

70. Lê et al., FactoMineR:an R package for multivariate analysis. J. Stat. Soft (2008).

71. Kolde et al. (2018).

